# Double Mutations in *Plasmodium falciparum* Kelch13 drive resistance to next-generation artemisinin derivatives in malaria parasites

**DOI:** 10.64898/2026.04.02.716214

**Authors:** Christopher Bower-Lepts, Kurt E. Ward, Sergio Wittlin, Barbara H. Stokes, Tomas Yeo, Tarrick Qahash, Jennifer L. Small-Saunders, Heekuk Park, Anne-Catrin Uhlemann, Manuel Llinás, David A. Fidock, Sachel Mok

## Abstract

New antimalarial compounds are urgently required to overcome artemisinin partial resistance that has emerged in Asia and now Africa. Ozonides are promising next-generation artemisinins that offer the improved pharmacokinetic property of a prolonged *in vivo* half-life. To assess the potential for parasite resistance to ozonides in an artemisinin-resistant background, we subjected Cambodian Kelch13 (K13) mutant parasites to increasing artefenomel (OZ439) pressure up to *in vivo* physiological concentrations. Whole-genome sequencing identified a novel non-propeller K13 A212T mutation in OZ439-resistant parasites. Gene editing and drug susceptibility assays revealed that the K13 double mutation R539T+A212T is a determinant of OZ439 resistance. In extended parasite recovery assays, this resistance mechanism was associated with accelerated parasite recrudescence following OZ439 or OZ277 exposure. This phenotype was also observed in K13 C580Y+A212T double mutant parasites. Global metabolomic profiling revealed no changes in the levels of hemoglobin-derived peptides in OZ439-resistant parasites, suggesting that resistance is not associated with drug activation. Instead, double mutant parasites exhibited increased levels of metabolites linked to glutathione, nucleotide, and aspartate-glutamate metabolism, suggesting a higher capacity for redox regulation to tolerate drug-induced oxidative damage. Our findings demonstrate that ozonide resistance can emerge through a novel K13 mutation on the background of existing artemisinin-resistance *k13* alleles.

## Introduction

Malaria remains a major global health challenge, causing an estimated 282 million cases and 610,000 deaths in 2024, the vast majority attributable to *Plasmodium falciparum* infection (*1*). Chemotherapeutic treatment of clinical malaria with effective antimalarial medicines remains critical to control efforts. The management of uncomplicated *P. falciparum* malaria relies primarily on artemisinin-based combination therapies (ACTs), which pair a swift-acting artemisinin derivative (dihydroartemisinin, artesunate, or artemether) with a longer-acting partner drug to achieve rapid parasite clearance and eliminate residual parasites (*2*).

Antimalarial drug resistance has limited progress in reducing the global malaria burden (*3*). Of particular concern is the emergence of artemisinin partial resistance, defined as delayed parasite clearance (clearance half-life ≥ 5 hours or day 3 parasite positivity following clinical treatment) or elevated survival (> 1%) in *in vitro* ring-stage survival assays (RSA) (*4*). First documented in Cambodia in the late 2000s, artemisinin partial resistance subsequently spread throughout Southeast Asia including Myanmar, Vietnam and Laos (*5–7*). Mutations in the propeller domain of the Kelch13 (K13) protein were identified as the genetic determinant of resistance using *in vitro* resistance selection, genome-wide association studies and gene editing approaches (*8–10*). The WHO now recognizes 13 mutations as validated markers of artemisinin resistance (ART-R), including C580Y, R539T and F446I that are predominant in Southeast Asia (*11*). Alarmingly, the independent emergence of resistance-associated K13 mutations linked to delayed parasite clearance has now been documented in East Africa and Horn of Africa, which bear a disproportionate burden of global malaria, with high rates of morbidity and mortality (*12–15*). In the context of emerging partner drug resistance, further erosion of ACT efficacy risks a devastating reversal in malaria control efforts.

Artemisinin derivatives are activated by heme-mediated cleavage of their endoperoxide bridge, producing carbon-centered radicals that alkylate a broad spectrum of parasite biomolecules inducing proteotoxic and oxidative stress resulting in cellular dysfunction and parasite death (*16–20*). Mutations in K13 lead to reduced drug activation by lowering hemoglobin endocytosis and limiting the levels of hemoglobin-derived free heme required for activation of artemisinins (*21*, *22*). Aside from lowered drug activation, K13 mutations are also associated with enhanced cellular stress tolerance, including increased protein turnover, modulation of the unfolded protein response, increased ubiquitin-proteasome activity and improved antioxidant defences (*23–26*). Early ring-stage K13 mutant parasites are able to survive the short *in vivo* half-life (∼1-hour) of artemisinin derivatives by entering quiescent, stress-adapted states and later resuming growth once drug concentrations decline.

Ozonides (1,2,4-trioxolanes) were developed as next-generation peroxide antimalarials intended to overcome key limitations of artemisinin derivates, including their short *in vivo* half-life, dependence on plant-based production, and the risk of emerging ART-R (*27*). Ozonides are fully synthetic and chemically stable, while retaining the essential endoperoxide pharmacophore required for antimalarial activity (*28*, *29*). Accordingly, functional studies and chemoproteomics have demonstrated substantial overlap in the mechanism of action of artemisinin and ozonide compounds, consistent with a shared heme-dependent activation and overlapping target profile (*18*, *30–32*). In contrast to artemisinins, the optimized trioxolane scaffold in ozonide compounds confers a substantially prolonged *in vivo* half-life of ∼42 hours, increasing systemic exposure and supporting the prospect of a single-dose therapy (*33*).

OZ439 (artefenomel) was engineered with enhanced steric stabilization of the peroxide bond to extend systemic exposure and reduce clearance relative to the first generation ozonide OZ277 (arterolane) (*34–38*). OZ439 exhibits rapid parasiticidal activity, high oral bioavailability, and activity across all asexual blood stages of *P. falciparum*, with similar pharmacokinetic profiles in malaria patients and uninfected volunteers (*34*, *39*, *40*). Importantly, *in vitro* studies have demonstrated OZ439 activity against parasites harboring *K13* mutations (*41–43*). OZ439 has progressed through Phase I and II clinical trials, initially in combination with piperaquine and more recently with ferroquine, though development has been constrained by formulation challenges (*44–46*). Nonetheless, recent studies describing novel trioxolane chemotypes with improved metabolic stability, favorable pharmacokinetic properties, and potent activity against artemisinin-resistant parasites highlight the continued promise of ozonide scaffolds for next-generation antimalarial development (*47*, *48*).

Despite the advanced clinical development of OZ439, the genetic basis and evolutionary trajectory of *P. falciparum* resistance to OZ439 has not been explored. In this study, we sought to assess the potential of *P. falciparum* to generate resistance to OZ439. Using *in vitro* resistance selection and whole-genome sequencing we identified the genetic determinant of *P. falciparum* resistance to ozonides. By employing gene editing, comprehensive phenotypic characterization and metabolomic profiling, we elucidated the molecular mechanisms of OZ439 resistance.

## Results

### Long-term *in vitro* selection with OZ439 identified a novel K13 A212T mutation

To identify genetic determinants of ozonide resistance, we subjected Cambodian Cam3.II *P. falciparum* parasites possessing a K13 R539T mutation to prolonged *in vitro* drug selection using OZ439 (Fig. 1). Cam3.II was selected as a clinically relevant Southeast Asian genetic background which has recently expanded in Laos (*49*).

**Fig. 1.**
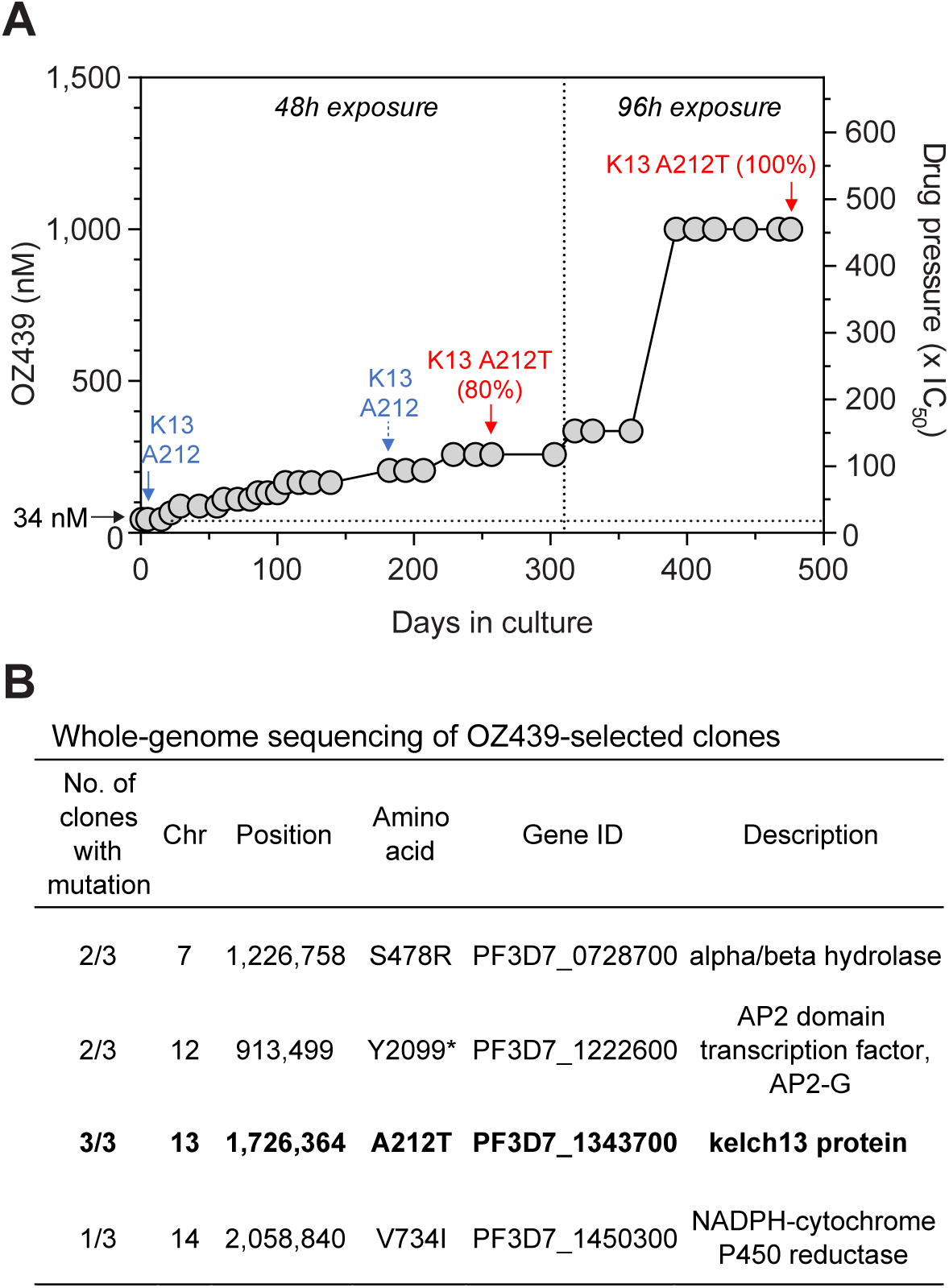
*In vitro* resistance selection and whole-genome sequencing identified a novel K13 A212T mutation as a putative molecular determinant of OZ439 resistance. (A) Cam3.II parasites were treated with increasing concentrations of OZ439 from 44 nM up to 1 µM in 48- or 96-hour exposures for a total of 33 drug-selection cycles to generate resistant parasites over 476 days in culture. Parasite lines were harvested for whole-genome sequencing (WGS) at selected timepoints (solid arrows) or Sanger sequencing (dashed arrow) at the intermediate timepoint. Blue arrows denote the detection of the wild-type K13 A212 sequence while red arrows denote the detection of the mutant K13 A212T sequence. The clinically relevant *in vitro* OZ439 concentration (34 nM) corresponding to a 500 mg dose in patients is annotated on the left Y-axis. **(B)** List of non-synonymous mutations identified in the OZ439-resistant clones from comparative WGS analysis against the OZ439-sensitive parent. These clones were obtained by limiting dilution of the final resistance-selected bulk population. Only the K13 A212T mutation was present in all three clones.

To generate OZ439 resistance *in vitro,* highly synchronous ring-stage Cam3.II parasites were exposed to escalating concentrations of OZ439, beginning at 44 nM (20x IC_50_) and incrementally increasing to 1 μM (∼450x IC_50_) (Fig. 1A). Drug pressure was initially applied in 48-hour pulses and later extended to 96-hour exposures for a total of 33 drug selection cycles, encompassing 476 days in culture. From day 359 onwards, parasites were able to proliferate under continuous exposure to 1 μM OZ439, indicating the acquisition of high-level resistance in the parasite population (Fig. 1A).

To identify the genetic changes associated with resistance, parasites were cloned by limiting dilution at the assay endpoint and independent clones were subjected to whole-genome sequencing (WGS) (Fig. 1B). Parental and OZ439-selected lines sequence reads yielded a depth of 31-42ξ coverage and with at least 93% of the genome having 10 or more reads. Comparative WGS analysis identified a shared nonsynonymous A212T mutation in K13 present in all clones sequenced at day 476 (15.5 months) (Fig. 1B). Sequencing at earlier timepoints revealed that this mutation was detected at 80% prevalence at day 257 (8.5 months) but was absent at day 182 (6 months) post-selection initiation. Additional mutations were detected in other clones, though none were detected in all clones, suggesting that these were not primary drivers of resistance. The fixation of the K13 A212T mutation in the parasite population under selection suggests that it is a putative driver of OZ439 resistance.

### K13 A212T mutants did not alter OZ439 susceptibility but retained resistance to DHA when combined with R539T or C580Y in RSAs

To characterize the contribution of K13 A212T - alone and in combination with the pre-existing R539T mutation or with K13 C580Y - to parasite susceptibility to OZ439, we used CRISPR/Cas9 genome editing to insert or remove the A212T mutation from parental Cam3.II parasites and resistant clones generated by *in vitro* selection (fig. S1). We established a panel of parasite lines comprising single (A212T, R539T or C580Y) and double (R539T+A212T or C580Y+A212T) K13 mutant lines together with an isogenic wild-type K13 control (Table 1 and fig. S1) and then assessed *in vitro* drug susceptibility of parasites to OZ439 and DHA (Fig. 2 and fig. S2).

**Fig. 2.**
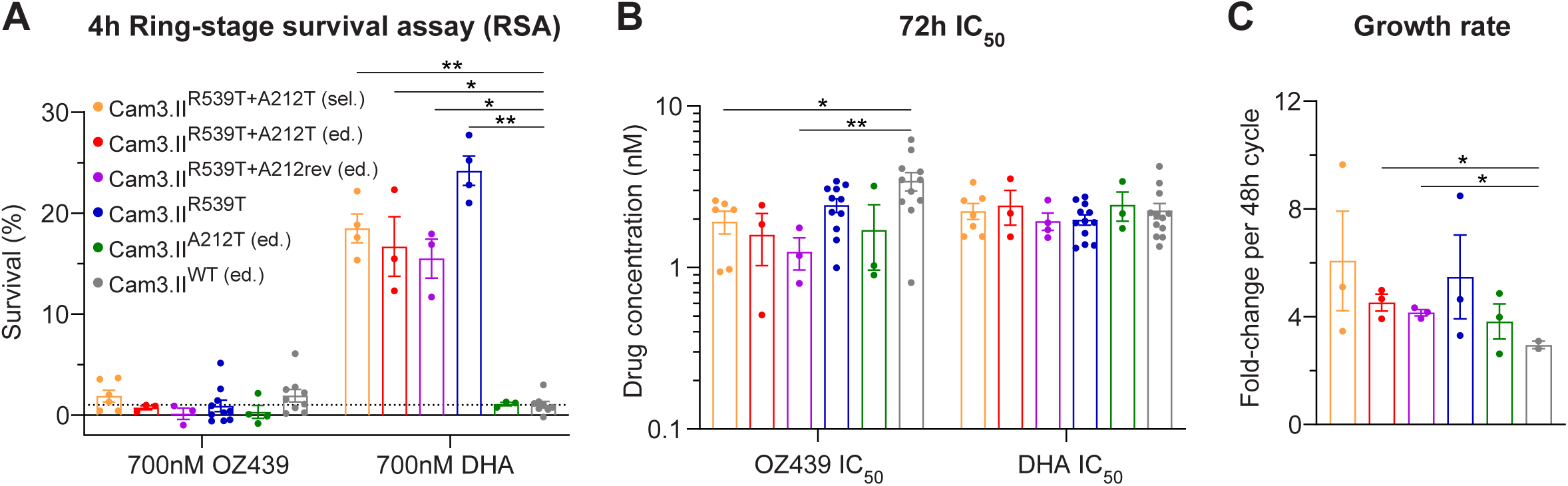
K13 R539T+A212T double mutant parasites retained resistance to DHA but did not exhibit altered responses to OZ439 in 4-hour or 72-hour drug susceptibility assays. **(A)** Percentage ring-stage survival of OZ439-selected, *k13* gene-edited and parental Cam3.II parasite lines following 4-hour exposure to 700 nM OZ439 or DHA. The dashed line at 1% survival denotes the threshold for *in vitro* ART resistance in ring-stage survival assays. **(B)** IC_50_ values of OZ439 and DHA against parasite lines determined from 72-hour dose-response curves. **(C)** Growth rates of parasite lines expressed as fold-change per 48-hour cycle corresponding to parasite replication rate. Bars in each graph are colored according to the corresponding parasite line displayed in the panel **A** legend. Error bars in each graph represent the standard error of the mean (SEM). Statistical significance was determined between isogenic wild-type and K13 mutant lines using an unpaired Welch’s t-test and significance is annotated above bars (*P < 0.05, **P < 0.01). Absence of annotation indicates no statistically significant difference was observed.

**Table 1.** Phenotype and genotype information for the panel of *k13* gene-edited parasite lines studied herein.

| Parasite lines | Drug used in selection | Edited in this study | *k13* allele | Secondary mutations | DHA phenotype | OZ439 phenotype |
| --- | --- | --- | --- | --- | --- | --- |
| Cam3.II<sup>R539T+A212T</sup> (sel.) | OZ439 | No | R539T+A212T | Yes<sup>‡</sup> | Resistant | Resistant |
| Cam3.II<sup>R539T+A212rev</sup> (ed.) | - | Yes | R539T | Yes<sup>‡</sup> | Resistant | Sensitive |
| Cam3.II<sup>R539T</sup> | - | No | R539T | - | Resistant | Sensitive |
| Cam3.II<sup>R539T+A212T</sup> (ed.) | - | Yes | R539T+A212T | - | Resistant | Resistant |
| Cam3.II<sup>WT</sup> | - | No<sup>†</sup> | WT | - | Sensitive | Sensitive |
| Cam3.II<sup>A212T</sup> (ed.) | - | Yes | A212T | - | Sensitive | Sensitive |
| Cam3.II<sup>C580Y</sup> | - | No<sup>†</sup> | C580Y | - | Resistant | Sensitive |
| Cam3.II<sup>C580Y+A212T</sup> (ed.) | - | Yes | C580Y+A212T | - | Resistant | Resistant |

<sup>†</sup>These two parasite lines were previously generated from the Cam3.II<sup>R539T</sup> parental line by editing the *k13* gene using zinc-finger nuclease-based editing.
<sup>‡</sup>Secondary mutations present in the OZ439-selected Cam3.II line comprise a S478R mutation in alpha/beta hydrolase (PF3D7\_0728700) and a Y2099\* nonsense mutation in AP2-G (PF3D7\_1222600)
sel.: selected; ed.: edited

In 4-hour ring-stage survival assays (RSA) using 700 nM OZ439, all lines remained sensitive to the drug, indicating no measurable survival advantage in double or single mutants relative to controls (Fig. 2A, fig. S2A). In contrast, all mutant parasites harboring R539T or C580Y displayed significantly elevated survival following 700 nM DHA exposure relative to sensitive controls (ranging between 10-25% survival) indicating that addition of A212T did not reduce survival of early rings to DHA. A212T alone did not confer increased DHA survival above the 1% threshold, suggesting that it does not independently drive ART-R (Fig. 2A).

To further examine whether the A212T mutation drives parasite resistance across other stages of the asexual cycle, we performed standard 72-hour IC_50_ assays using OZ439 and DHA. All lines profiled in the assay exhibited comparable sensitivity to OZ439 (Fig. 2B, fig. S2B). Similarly, none of the single or double K13 mutant lines displayed reduced sensitivity to DHA, including those possessing R539T or C580Y mutations, which are linked only to early ring-stage survival (Fig. 2B, fig. S2B). Together, these findings indicated that the K13 A212T mutation does not confer reduced susceptibility to OZ439 in either RSAs or conventional 72-hour IC_50_ assays, either alone or in combination with the DHA resistance-conferring K13 mutations R539T or C580Y.

### The K13 R539T+A212T and C580Y+A212T double mutants exhibited accelerated recovery following OZ439 or OZ277 exposure

To characterize the survival phenotype in the OZ439-selected parasites, we examined how parasites recovered following exposure to physiologically relevant concentrations of drug. Parasites from the panel of *k13* gene-edited and drug-selected lines were exposed to a range of OZ439 concentrations (up to 1 μM) for 48 hours and parasitemia was monitored by flow cytometry every 48-72 hours for up to 14 days (Fig. 3 and fig. S3).

**Fig. 3.**
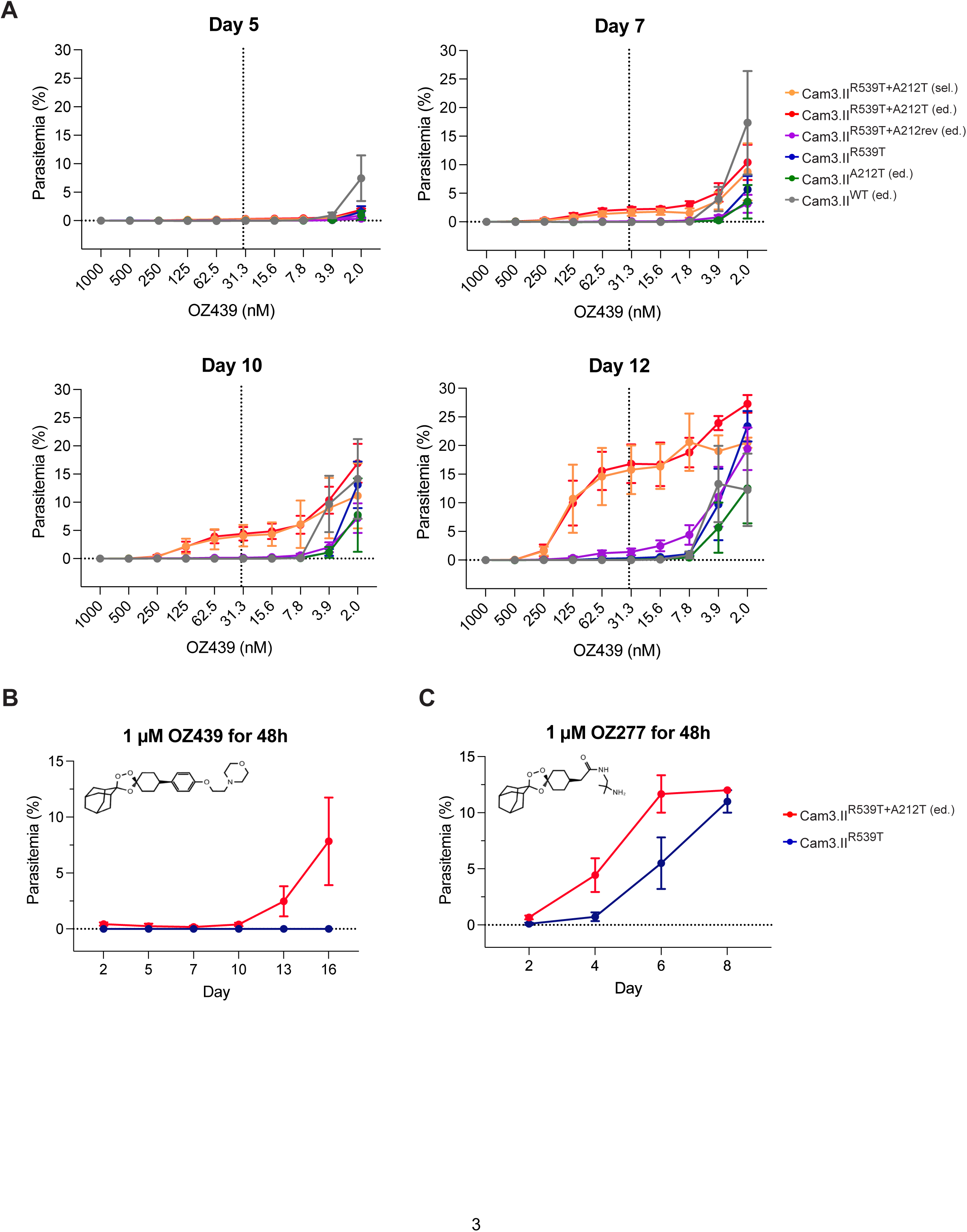
K13 R539T+A212T parasites exhibited enhanced survival and recovery following 48-hour exposure to OZ439 and OZ277. **(A)** Parasitemia of OZ439-selected, *k13* gene-edited and parental Cam3.II parasite lines following exposure to serial dilutions of OZ439 for 48 hours measured at day 5-12 following initiation of drug treatment as determined by flow cytometry. The vertical dashed line in each graph denotes the clinically relevant OZ439 concentration of 34 nM. (**B** and **C**) Parasitemia of Cam3.II^R539T+A212T^ and Cam3.II^R539T^ lines measured for up to 14 days following 48-hour exposure to 1 μM OZ439 (B) or OZ277 (C) determined by microscopy. Chemical structures for OZ439 and OZ277 are shown in upper left of graphs. Points in each graph represent mean values determined across three independent experiments with technical duplicates. Error bars in each graph represent the SEM. Extended results for C580Y+A212T parasites are found in fig. S2.

No substantial differences in parasitemia were evident between lines at day 5 (Fig. 3A). However, from day 7 onwards viable parasites were detected for the R539T+A212T double mutants but not in single mutants and wild-type controls after treatment with 7.8 nM to 125 nM OZ439. (Fig. 3A). By day 12, double mutant parasites recovered robustly, displaying a markedly higher parasitemia than all other lines at and above clinically relevant drug concentrations (34 nM), equivalent to a 500 mg *in vivo* dose (*43*). In contrast, R539T and A212T single mutants, in addition to K13 wild-type lines, recrudesced only at the lower drug concentrations (3.9 nM and below) (Fig. 3A and fig. S3).

The C580Y+A212T double mutant demonstrated accelerated recovery relative to the C580Y single mutant following 48-hour exposure to a clinically relevant concentration of OZ439 (fig. S2C). Similarly, measurement of day 7 parasitemia across a range of OZ439 concentrations confirmed enhanced survival of the double mutant (fig. S2D), paralleling the recrudescence phenotype observed in the R539T+A212T parasites. To investigate whether the accelerated rate of recovery in double mutants was linked to intrinsic fitness advantages, we measured parasite replication rates in the absence of drug. No differences in growth rates between single and double mutants were observed, suggesting that OZ439 resistance is linked to enhanced drug tolerance and faster recovery (Fig. 2C, fig. S2E).

To validate this phenotype using a different parasite quantification method, we challenged the gene-edited R539T+A212T double mutant line and the parental R539T single mutant with a single high dose of 1 μM OZ439 for 48 hours and monitored parasite growth by microscopy (Fig. 3B). Parasites in the double mutant line were detectable by day 10 and parasitemia exceeded 5% by day 16, in contrast to the R539T single mutant which failed to recover. Treatment with the first-generation ozonide OZ277 also resulted in accelerated recovery in the double mutant, indicating that the R539T+A212T genotype may mediate cross-resistance within the ozonide class of compounds (Fig. 3C). These findings confirm that addition of A212T to the R539T background confers a distinct survival phenotype following OZ439 exposure, which manifests as accelerated post-treatment recovery following drug exposure.

### K13 R539T+A212T double mutants survived physiologically relevant OZ439 exposure *in vivo*

To determine whether the K13 R539T+A212T genotype also conferred reduced susceptibility to OZ439 under *in vivo* conditions, we next evaluated the therapeutic efficacy of OZ439 using a humanized mouse model (Fig. 4). NOD SCID Gamma (NODscidIL2Rγ^null^) mice were engrafted with human erythrocytes and infected with either CRISPR-edited R539T+A212T double mutant parasites, R539T single mutants or isogenic wild-type controls. On day 3 post-infection, mice received a single dose of 10 mg/kg OZ439(*50*). Parasitemia was monitored for up to 24 days by both microscopy and flow cytometry to assess recrudescence dynamics (Fig. 4A and fig. S4).

**Fig. 4.**
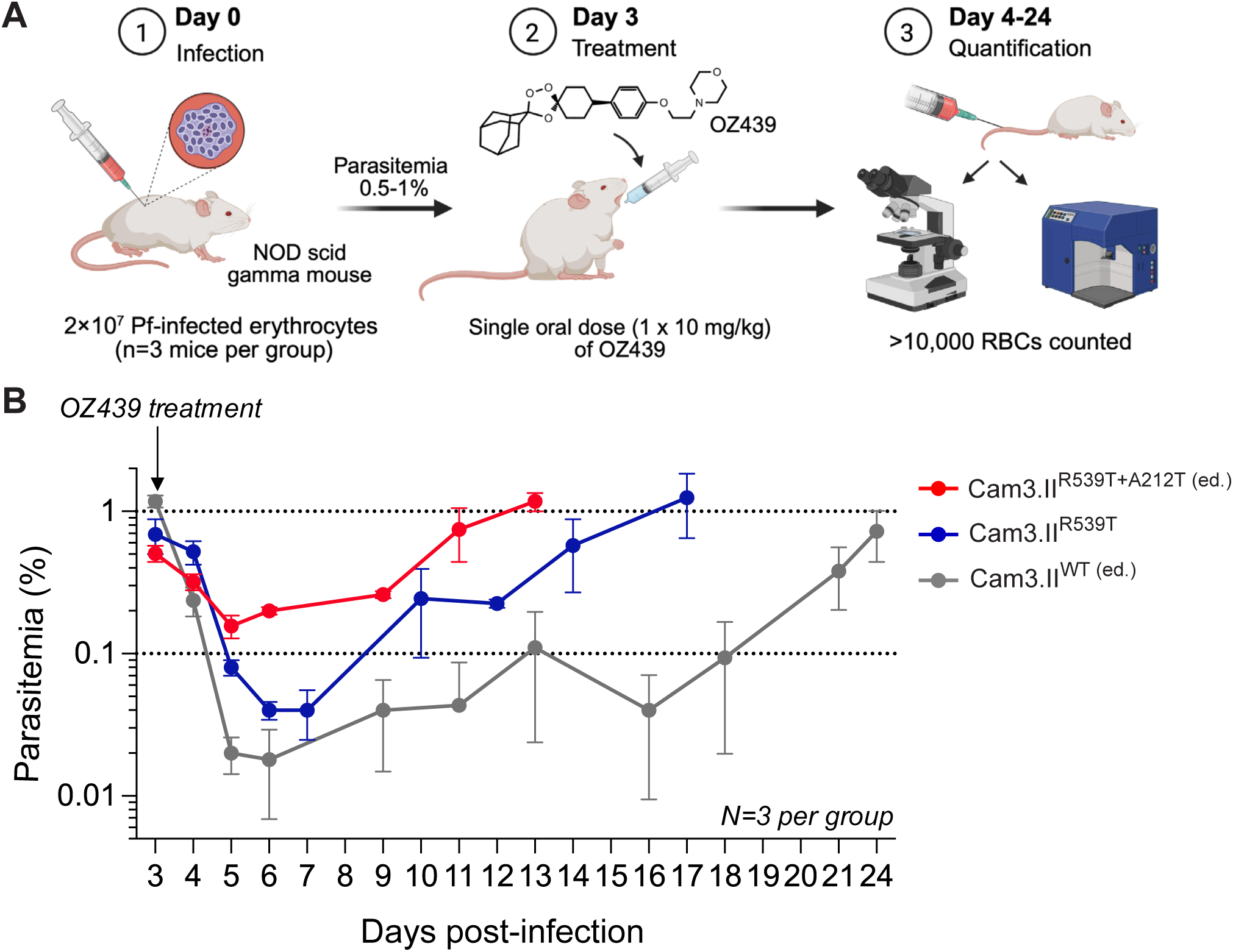
The K13 R539T+A212T double mutation conferred *in vivo* resistance to physiologically relevant concentrations of OZ439. **(A)** Schematic overview of workflow for evaluation of the *in vivo* efficacy of OZ439 using the NODscidIL2Rγ^null^ (NSG) mouse model. Three mice per parasite strain were engrafted with 2×10^7^ human erythrocytes infected with *in vivo-*adapted Cam3.II^R539T+A212T^, Cam3.II^R539T^ or Cam3.II^WT^ parasites. A single oral dose of 10 mg/kg OZ439 mesylate was administered once parasitemia reached 0.5-1%. From day 4 onwards, parasitemia was measured by microscopic examination of Giemsa-stained blood smears or flow cytometry every 1-3 days for up to 24 days. Images were created with BioRender.com. **(B)** Parasitemia of Cam3.II^R539T+A212T^, Cam3.II^R539T^ and Cam3.II^WT^ lines over 21 days following administration of OZ439 mesylate to mice (depicted by the arrow). Values represent the mean parasitemia in the peripheral blood of three individual mice obtained from microscopic counts of Giemsa smears. Error bars represent the SEM.

Consistent with the recovery phenotype observed following *in vitro* exposure, the R539T+A212T double mutant exhibited substantially accelerated recrudescence relative to single mutant parasites and controls, reaching 1% parasitemia by day 13 post-infection, compared to day 17 and day 24 for the R539T single mutant line and wild-type line respectively (Fig. 4B). These findings demonstrate that the addition of A212T to the R539T background confers a survival advantage under physiologically relevant OZ439 exposure *in vivo*, recapitulating the enhanced post-treatment recovery phenotype observed *in vitro*.

### Enhanced OZ439 recovery in K13 double mutant parasites did not associate with altered hemoglobin-derived peptide levels

We next sought to define the mechanism underpinning the post-OZ439 treatment recovery phenotype observed in K13 double mutant parasites. ART-R–associated K13 mutations, including R539T, have been shown to reduce K13 protein stability and impair hemoglobin endocytosis and digestion, which limits the release of heme required for activation of artemisinin derivatives (*21*, *22*, *25*). As OZ439 and artemisinins share structural and mechanistic features, we hypothesized that enhanced ozonide survival in K13 double mutant parasites might similarly arise from further inhibition of hemoglobin uptake or catabolism.

To test this, we performed stage-specific untargeted metabolomics in absence or presence of OZ439 and DHA for the panel of gene-edited K13 mutant and wild-type isogenic lines to quantify the levels of host hemoglobin-derived peptides (Fig. 5A). We first examined peptide abundance for each K13 edited line relative to the R539T single mutant line in the absence of drug treatment. As expected, WT parasites exhibited elevated levels of all peptide classes relative to the Cam3.II R539T line, consistent with the previous observation of impaired hemoglobin digestion associated with this mutation (Fig 5B to D). A212T single mutants exhibited peptide profiles comparable to wild-type parasites across all peptide classes. Similarly, the addition of A212T to R539T in double mutant parasites did not confer additional reductions in peptide levels in both trophozoites and rings (Fig. 5B to D). These findings suggest that the acquisition of A212T does not affect the levels of hemoglobin-derived peptides, thus may not impair hemoglobin digestion and downstream drug activation.

**Fig. 5.**
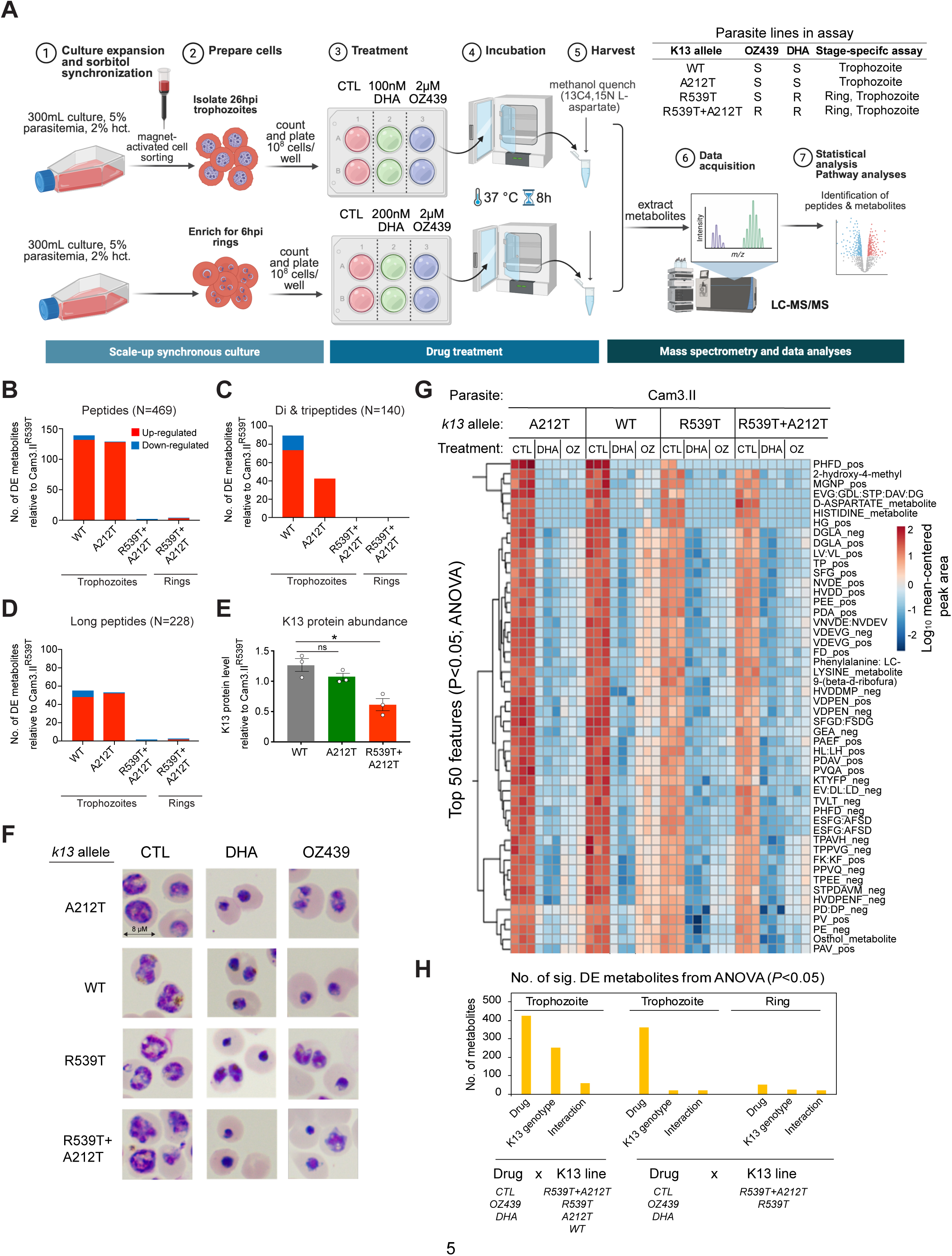
The K13 R539T+A212T mutation was not associated with altered levels of hemoglobin-derived peptides at basal level while OZ439 treatment caused downregulation of overall peptide levels irrespective of K13 genotype. **(A)** Schematic workflow of global metabolomic profiling experiment. The table lists the isogenic Cam3.II lines carrying various *k13* alleles used in this assay and their relevant phenotype and genotype information. ‘S’ denotes sensitive, and ‘R’ denotes resistant. Images were created with BioRender.com. Further details are available in Methods section. (**B** to **D**) Number of differentially expressed (DE) peptides (B), di- and tripeptides and (C) long peptides (D) in each K13-edited line compared against the Cam3.II^R539T^ line, as determined from pairwise t-tests. The total number of peptides analyzed within each category is annotated above graphs. Blue and red represent up- or down-regulated peptides, respectively. **(E)** K13 protein levels measured in K13-edited trophozoite stages lines using immunoblotting assays. Total K13 protein abundance including processed products was normalized to ERD2 protein loading controls in each blot and is expressed relative to Cam3.II^R539T^ parasites. Error bars represent the SEM. Statistical significance was determined using unpaired Welch’s t-tests and is annotated above bars (*P < 0.05, ‘ns’ represents no statistical significance observed). **(F)** Brightfield images of K13-edited and parental Cam3.II parasite lines following treatment of trophozoites with 2 µM OZ439 or 100 nM DHA, or no-drug control for 8 hours**. (G)** Differences in peptide levels in isogenic K13-edited lines following treatment with OZ439 or DHA or no-drug control in trophozoite stage. Top 50 features were determined using ANOVA (P<0.05). For ring-stage treatments refer to fig. S5A. **(H)** The number of differentially expressed metabolites detected in three separate analyses comparing; i) the effect of drug treatment (OZ439, DHA and no-drug control) and *k13* genotypes (R539T+A212T, R539T, A212T, Wild-type) in trophozoite stage, ii). the effect of drug treatment (OZ439 and no-drug control) and *k13* genotypes (R539T+A212T and R539T) in trophozoite and ring stages. Significance was determined by ANOVA (P<0.05).

To determine whether altered K13 protein abundance might explain these findings, we quantified K13 levels in trophozoites by immunoblotting assays (Fig. 5E). A212T single mutants displayed K13 protein levels comparable to wild-type parasites, whereas R539T+A212T double mutants exhibited a further reduction in K13 protein relative to R539T alone (Fig. 5E). Despite this additional decrease in K13 abundance, peptide levels were not further decreased in the double mutant, suggesting that impairment of hemoglobin digestion may be maximal in R539T parasites, and is not proportionally exacerbated by additional K13 destabilization.

### OZ439 and DHA treatment downregulated peptide levels independently of K13 genotype

We next examined the effects of drug treatment on parasite cellular morphology and metabolite abundance across genotypes. Treatment of trophozoites with 100 nM DHA or 2 μM OZ439 for 8 hours induced a pyknotic, condensed morphology in all lines irrespective of K13 genotype, whereas treatment of rings produced no striking differences in morphology (Fig. 5F, fig. S5A). DHA or OZ439 treatment resulted in significant reductions in peptide abundance in the drug-resistant and sensitive lines (Fig. 5G, fig. S5B and C). Notably, the magnitude of peptide depletion following OZ439 or DHA exposure did not differ between R539T single mutant and R539T+A212T double mutant lines, implying that the acquisition of A212T does not impact peptide processing in response to ozonide or DHA treatment (Fig. 5G, fig. S5B and C). Analyses of differentially expressed metabolites confirmed that the largest differences in peptide levels were driven by drug exposure. Minimal differences in peptide levels were observed in the OZ439-resistant K13 double mutant parasites compared to K13 single mutant and control parasites in response to DHA or OZ439 in ring and trophozoite stages (Fig. 5G and H).

Taken together, these data demonstrate that although the R539T+A212T double mutation may further reduce K13 protein abundance, it does not confer additional impairment of hemoglobin-derived peptide production under basal or drug-treated conditions. The enhanced OZ439 recovery phenotype observed in double mutant parasites therefore cannot be explained by further suppression of hemoglobin endocytosis or digestion, thus likely does not invoke reduced drug activation. These findings indicate that ozonide resistance mediated by the R539T+A212T genotype likely involves a distinct cellular pathway.

### OZ439 resistance in K13 double mutant parasites associated with enhanced redox and nucleotide metabolism in OZ439 resistance

To elucidate other possible cellular pathways associated with OZ439 resistance, we analyzed the metabolomic profiles in gene-edited R539T+A212T double mutant parasites to identify metabolites and pathways attributable to A212T. Metabolite levels in untreated ring- and trophozoite-stage parasites were quantified in the OZ439-resistant R539T+A212T double mutant parasites and compared to sensitive isogenic R539T single mutants (Data S1).

In ring-stage parasites, glutathione metabolism was significantly enriched in the double mutant relative to the R539T line (Fig. 6A and fig. S6). These parasites exhibited elevated levels of glutathione disulfide and cysteinylglycine (Cys-Gly), indicative of increased glutathione turnover and redox cycling (Fig. 6B). Pyrimidine metabolism was also strongly enriched in ring-stage parasites, with elevated levels of cytidine monophosphate (CMP) and N-carbamoyl-L-aspartate, suggesting enhanced nucleotide biosynthesis (Fig. 6B).

**Fig. 6.**
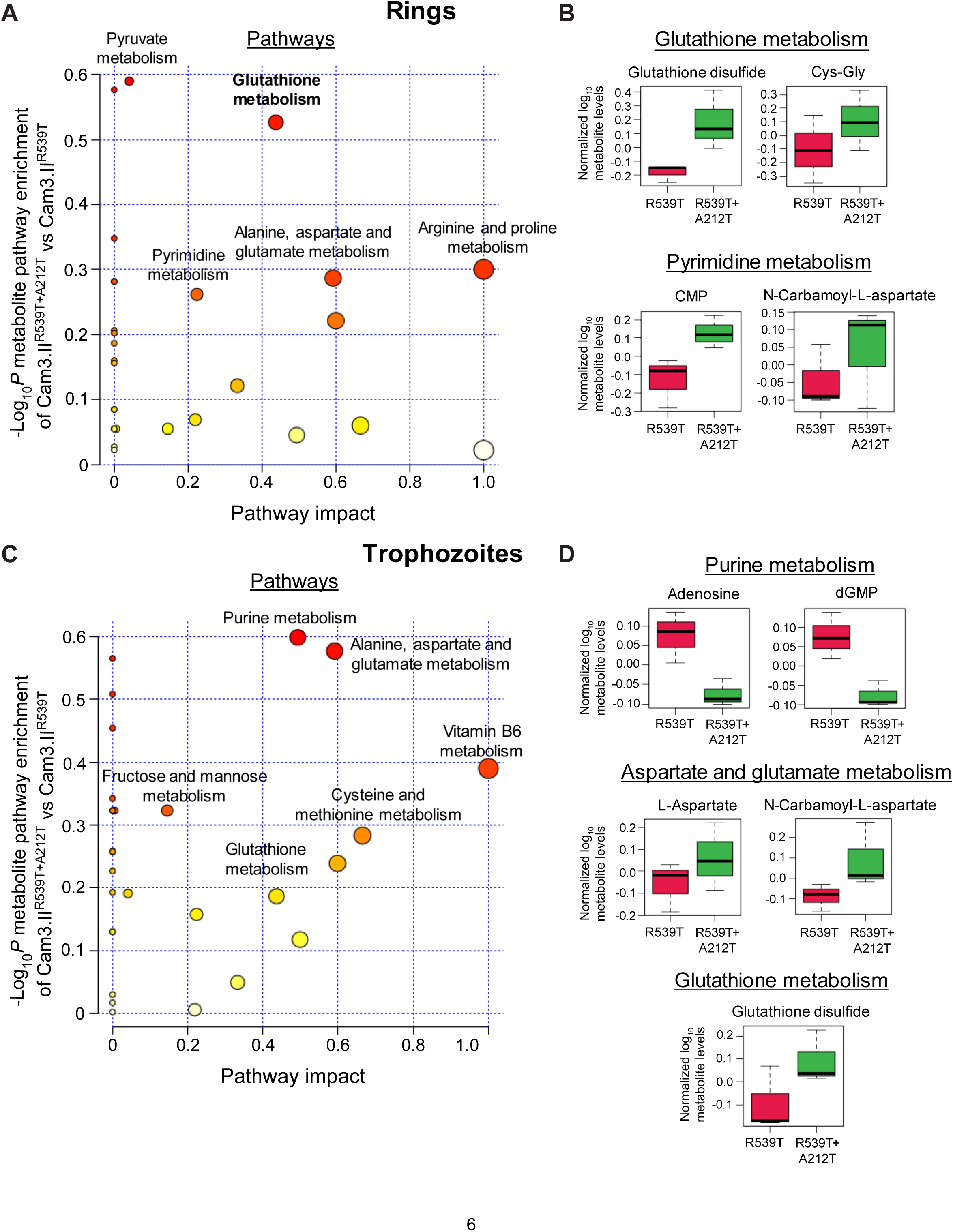
The K13 R539T+A212T double mutant displayed an increase in glutathione, pyrimidine and alanine, aspartate and glutamate metabolites in rings or trophozoites. **(A)** Metabolic pathway enrichment analysis of Cam3.II^R539T+A212T^ relative to Cam3.II^R539T^ in ring-stage parasites in the absence of drug treatment. **(B)** Normalized log_2_-transformed levels of glutathione and pyrimidine metabolites in Cam3.II^R539T^ and Cam3.II^R539T+A212T^ lines in ring stages. **(C)** Metabolic pathway enrichment analysis in Cam3.II^R539T+A212T^ relative to Cam3.II^R539T^ in trophozoite stage. **(D)** Normalized log_2_-transformed levels of purine, aspartate and glutamate, and glutathione metabolites in Cam3.II^R539T^ and Cam3.II^R539T+A212T^ parasite lines in the trophozoite stage. Metabolic pathways annotated in A and C are those displaying the highest -log_10_ *P* score in Cam3.II^R539T+A212T^. Circle size represents pathway impact value and are colored according to the enrichment -log_10_ *P* score.

In trophozoites we found convergence of several metabolic pathways enriched in ring stages, including glutathione, aspartate and glutamate metabolism. Elevated glutathione disulfide was detected in the double mutant as were increased levels of L-aspartate and N-carbamoyl-aspartate (Fig. 6C and D, fig. S7). In addition, purine metabolism was downregulated in R539T+A212T trophozoites compared to R539T parasites, revealed by reduced levels of adenosine and dGMP (Fig. 6C and D, fig. S7). In contrast, none of these pathways were enriched in K13 A212T single mutant parasites relative to wild-type controls, indicating that these metabolic changes were specifically associated with the double mutation in the OZ439-resistant parasites (fig. S8). Taken together, the enrichment of these metabolites in OZ439-resistant parasites suggests an increased engagement of antioxidant pathways across developmental stages and rewiring of key metabolites that interface with both nucleotide synthesis and redox balance.

## Discussion

In this study, we performed long-term *in vitro* selection of *P. falciparum* Cam3.II K13 R539T parasites under OZ439 pressure to investigate the capacity for ozonide resistance to emerge in artemisinin resistant parasites. Using whole-genome sequencing analysis, we identified a novel K13 A212T mutation, located in the non-propeller region of the protein, as a genetic determinant of resistance. Gene-editing and phenotypic analyses demonstrated that the R539T+A212T double mutation confers a distinct drug resistance phenotype characterized by accelerated parasite recovery following drug exposure and removal. This phenotype was observed both *in vitro* and *in vivo*, conferred cross-resistance to the related ozonide OZ277 and was recapitulated when A212T was introduced onto a K13 C580Y mutant background. Metabolomic profiling demonstrated that A212T-driven resistance was not associated with reduced levels of hemoglobin-derived peptides suggesting no alteration in rates of hemoglobin digestion, the primary described mechanism of ART-R. Instead OZ439 resistance associated with a distinct metabolic signature encompassing elevated levels of glutathione and pyrimidine metabolism.

These findings provide, to our knowledge, the first confirmed evidence of genetically defined resistance to an ozonide antimalarial emerging under sustained drug pressure. We identified K13 A212T as the first resistance-associated mutation upstream of the BTB/POZ and propeller domains, and validated it as a genetic determinant of antimalarial resistance. A212T is a novel K13 mutation which has not previously been detected in clinical isolates (*51*). Importantly, ozonide resistance manifests as an enhanced recovery phenotype that is not captured by conventional RSA or IC_50_ readouts where parasite survival is measured after one asexual replication cycle, highlighting a key blind spot in current assays. Instead, OZ439-resistant parasites exhibit delayed recrudescence following drug exposure, suggesting a capacity for these parasites to enter a dormant, drug-tolerant state and subsequently resume proliferation once drug pressure is removed. Similar metabolic adaptations leading to quiescent states with enhanced drug resilience have previously been described in artemisinin-resistant parasites (*25*, *52*). The recovery dynamics observed in OZ439-resistant parasites suggests that increased tolerance to endoperoxides may invoke a similar mechanism of dormancy. Mechanistically, the absence of hemoglobin digestion perturbation distinguishes this phenotype from canonical K13-mediated ART-R and instead implicates metabolic rewiring and pathways linked to redox balance and nucleotide biosynthesis that support parasite persistence under ozonide pressure and enhanced recovery (*21*). Ozonide endoperoxides are activated by heme but have also been shown to directly alkylate heme within *P. falciparum*, a process proposed to contribute to their antimalarial activity (*53*, *54*). Given that there was no difference in hemoglobin catabolism between resistant double mutants and sensitive lines, it is unlikely that resistance is mediated through the levels of heme-drug adduct formation. It is likely that metabolic adaptations affecting redox balance may influence the parasite’s ability to tolerate the oxidative and proteotoxic stress associated with heme alkylation.

Previous work has demonstrated that ozonide treatment perturbs biochemical pathways beyond the digestive vacuole functions, including lipid biosynthesis, pyrimidine metabolism and aspartate/glutamate metabolism (*55*). In that report, exposure to ozonides in drug-sensitive parasites resulted in depletion of L-aspartate and N-carbamoyl-L-aspartate, possibly via drug-induced damage to proteins involved in the pyrimidine biosynthesis pathway (*55*). We observed elevated levels of L-aspartate and N-carbamoyl-L-aspartate in OZ439-resistant K13 double mutant parasites, which may reflect a compensatory mechanism for the loss of these cellular functions due to drug exposure. Our findings also detected elevated glutathione disulfide and Cys-Gly levels, indicative of increased glutathione turnover and enhanced redox cycling that may enable more efficient buffering of ozonide-induced oxidative stress, permitting parasites to withstand biomolecular damage and restore redox homeostasis following drug removal (*56*). Together, these metabolic changes may prime double mutant parasites to withstand oxidative insults generated by OZ439-induced proteotoxic stress and lipid peroxidation.

Metabolomic profiling under OZ439 exposure did not reveal any striking differences between resistant and sensitive parasites. One possibility is that the short drug exposure may not capture metabolic changes associated with enhanced recovery observed only from day 7 onwards. Recent work has shown that transcriptomic rewiring occurs from day 5 in dormant parasites which recrudesce following DHA treatment (*57*). Future gene expression studies incorporating extended sample collection timepoints will be useful to elucidate which pathways mediate accelerated recovery post-drug exposure in OZ439-resistant parasites. Similarly, the mechanistic link between the K13 A212T mutation and protein function remains unexplored. Unlike the propeller-domain mutations implicated in ART-R, A212T is located in the *Plasmodium*-specific region of the K13 protein. One possibility is that non-propeller substitutions modify protein-protein interactions, subcellular localization or post-translational regulation, thereby reshaping downstream signaling or stress response pathways. Defining the K13 interactome in basal conditions and in double mutant parasites will be essential to characterize genotype-phenotype associations, and to determine whether metabolic rewiring is a general mechanism of endoperoxide resistance or a consequence of specific K13 mutations.

The generation of OZ439 resistance described here raises important considerations for the future development and deployment of ozonide antimalarials. Although assessment of OZ439 in phase clinical trials has highlighted formulation challenges associated with this compound, substantial investment continues in next-generation ozonides and reformulated OZ439-adjacent chemotypes with improved pharmacokinetic and pharmacodynamic properties are being developed (*45–48*). Our findings demonstrate that high-level OZ439 resistance can emerge after ∼8 months of *in vitro* selection, indicating that while resistance is not biologically constrained, it arises only after prolonged selection, consistent with a high barrier to its evolution. Nonetheless, should optimized ozonides progress to widespread deployment, similar selective pressures could plausibly emerge in endemic settings. Of particular concern is the emergence of resistance on pre-existing K13 mutant backgrounds, as mutations in K13 are now well established in Southeast Asia and are increasing in prevalence in parts of Africa. Our data suggests that these genotypes may provide a permissive background for the acquisition of additional mutations, such as A212T, that could confer reduced ozonide susceptibility. Given that enhanced recovery in these OZ439-resistant parasites may translate to recrudescence or treatment failure in patients, more comprehensive drug susceptibility assays that extend the observation window or quantify recovery dynamics are required. The identification of a non-propeller domain mutation contributing to resistance underscores the need to broaden molecular surveillance beyond the propeller region of K13 in field samples. Early detection of the emergence of these mutations in parasite populations would be valuable to anticipate the potential impact on future ozonide deployment.

Collectively, this work establishes a novel genetic and biological route to ozonide resistance, identifying K13 A212T as a driver of reduced OZ439 susceptibility and implicating metabolic adaptation as the primary mechanism of parasite survival. These findings also expand the functional landscape of resistance-associated K13 mutations beyond the propeller domain and underscore the importance of expanding genomic surveillance to account for resistance phenotypes that extend beyond conventional susceptibility metrics. Anticipating and detecting such non-canonical resistance mechanisms will be critical to safeguarding the therapeutic lifespan of endoperoxide antimalarials and to informing rational strategies for their clinical deployment.

## Materials and Methods

### P. falciparum lines and in vitro culture

Asexual blood stage *P. falciparum* parasites were cultured *in vitro* at 3% hematocrit in human O^+^ RBCs (Interstate blood bank, USA) and in complete RPMI 1640 medium supplemented with 0.5% AlbuMAX II (Thermo Fisher Scientific). Cultures were maintained at 37°C and in 5% O_2_ and 5% CO_2_ as described (*25*). Parasite lines were genotyped by Sanger sequencing of the *k13* gene to verify their identities before the start of an experiment.

### *In vitro* selection of OZ439 resistance

OZ439 resistance was obtained by applying a ramping selection approach on the Cam3.II K13 R539T parasite line. A synchronized Cam3.II culture at 5% ring-stage parasitemia in 15 mL volume (10^8^ parasites) was exposed to OZ439 drug at 20× IC_50_ concentration for 48 hours. Following drug wash-off, parasites were allowed to recover in drug-free media until parasitemia reached 5% as monitored by microscopy. This selection cycle was repeated until the parasite recovery time shortened from ∼14 days to ∼3-5 days. Thereafter, in subsequent cycles, the drug concentration was increased stepwise by ∼30% starting from 44 nM until 1 µM and with exposure duration extended from 48-96 hours. The final drug-selected Cam3.II bulk was cloned by limiting dilution to generate clones that were whole-genome sequenced.

### *In vivo* therapeutic efficacy of OZ439 in NODscidIL2Rγ^null^ mouse model

*In vivo* adaptation of asexual blood stage *P. falciparum* lines, Cam3.II^R539T^ parent, Cam3.II^R539T+A212T^ and Cam3.II^WT^ was first achieved through three passages in NODscidIL2Rγ^null^ (NSG) mice performed over ∼3 months (*58*, *59*). The assessment of *in vivo* OZ439 efficacy against these three strains was tested by first infecting NSG mice engrafted with human RBC intravenously with 2×10^7^ parasitized RBCs per strain/mouse on day 0. Three mice were used per parasite strain. Once parasitemia reached 0.5-1% on day 3, the mice were treated with a single oral dose of 10 mg/kg of OZ439 mesylate compound (solubilized in 70% Tween-80 and 30% ethanol and diluted in water). Parasitemia was measured by microscopic examinations of Giemsa smears and flow cytometry every 1-3 days until day 24 by sampling 2 µL of blood from the mice at these time points.

### Whole-genome sequencing of OZ439-selected parasites

Parasite cultures at 1-3% parasitemia were lysed with 0.1% saponin in 1× PBS, and genomic DNA was extracted using the QIAamp DNA Blood Mini Kit (Qiagen). Libraries were prepared from 150 ng of genomic DNA using the Illumina DNA prep protocol. The quantified libraries were pooled and sequenced on an Illumina MiSeq using a 600-cycle kit (2 × 300 bp paired-end reads). Sequence data were aligned to the *P. falciparum* 3D7 genome (PlasmoDB version 36.0) and processed as described previously (*60*) using Genome Analysis Toolkit Haplotype Caller and snpEFF to call and annotate high quality variants, respectively. SNPs in the core genome that differed between the parental Cam3.II line and the OZ439-selected clones were identified by retaining only those that were homozygous, based on >90% alternate allele frequency and with at least 5 alternate reads.

### Drug susceptibility and growth assays

#### 4-hour ring-stage survival assays

Tightly-synchronized 0-3-hour early ring-stage parasites were obtained by incubating Percoll-density gradient purified schizonts with fresh RBCs for 3 hours followed by treatment with 5% D-Sorbitol, as described previously (*61*). Parasites seeded at 0.4% parasitemia and 1% hematocrit were exposed to 700 nM DHA or OZ439 for 4 hours at 37°C in 96-well plates. Thereafter, cultures were washed four times with complete medium and transferred to fresh 96-well plates using the Tecan Freedom EVO MCA96 instrument. Cultures were subsequently maintained for an additional 66 hours in drug-free medium. Assays were performed on three independent occasions with technical duplicates.

#### 72-hour drug susceptibility assays

Ring-stage parasites seeded at 0.4% parasitemia and 1% hematocrit were exposed to DHA and OZ439 across a 10-point drug dilution series alongside 0.1% DMSO (vehicle-treated) controls. Assays were performed on three independent occasions with technical duplicates.

#### 48-hour drug exposure assays

Ring-stage parasites were exposed to a 10-point dilution of OZ439 for 48 hours. Drug removal at 48-hour post-exposure was performed as described above. Cultures were replenished with media at 2 to 3-day intervals and fresh RBCs were added on day 7. Parasitemia was measured by flow cytometry every 2-3 days up to a 2-week period. Assays were performed on three independent occasions with technical duplicates.

#### Flow cytometry to quantify viable parasites

Parasitemia for drug-treated and vehicle control-treated wells were measured by flow cytometry, as described previously (*25*). Parasites were incubated with 1× SYBR Green (ThermoFisher) and 100 nM MitoTracker DeepRed (ThermoFisher) for 30 min at 37°C and quenched with 1× PBS. On average, 10,000 cells were analyzed per sample, using an iQue Screener Plus flow cytometer (Sartorius). Viable parasites were defined as the percentage of MitoTracker-positive and SYBR Green-positive labelled cells. For all drug assays, the percentage parasitemia for kill control wells (1 µM DHA) was subtracted from the parasitemia measured for each well.

#### Calculation of RSA and IC_50_ values

Parasite survival in the presence of DHA was expressed as the percentage of the background-subtracted parasitemia of the DHA or OZ439-treated samples divided by the DMSO-treated samples. Mean RSA survival rates >1% were defined as DHA resistant (*62*). The IC_50_ defines the drug concentration that results in 50% inhibition of parasite growth. 72-hour IC_50_ values were determined by applying a non-linear regression model on the normalized % survival across the log-transformed drug concentrations using Prism v8.3.1 (GraphPad). Unpaired Welch’s t-tests were performed using Prism v8.3.1 to determine if differences in RSA and IC_50_ values observed between lines were statistically significant.

#### Gene editing of *k13* A212T by CRISPR/Cas9

CRISPR/Cas9 editing was performed to introduce or remove the A212T mutation from the *k13* locus in Cam3.II clones, using the all-in-one plasmid pDC2-cam-coSpCas9-U6-gRNA-h*dhfr* as previously described (*63*), with the following modifications. Cloning of gRNAs was performed using primer pair p1+p2. A 760 bp donor template located within the *k13* gene was amplified from the parental Cam3.II line containing A212 or from the OZ439-selected Cam3.II line containing A212T using the primer pair p3+p4 and separately cloned into the pGEM T-easy vector system (Promega). For each donor construct containing A212 or A212T, we introduced 3 silent shield mutations at the Cas9 cleavage site to protect the edited locus from subsequent cleavage. Site-directed mutagenesis was performed on the donor sequences to introduce these silent mutations using the primer pair p5+p6. Donor templates were amplified using the primer pair p7+p8 and cloned into the final vector at the EcoRI and AatII restriction sites by In-Fusion Cloning (Takara) to generate the plasmids pDC2-cam-coSpCas9-U6-gRNA-K13_bsm-h*dhfr* and pDC2-cam-coSpCas9-U6-gRNA-K13_A212T-h*dhfr*. All final plasmids were verified by Sanger sequencing using primers p9 to p11. All primer sequences are listed in Table S1. To obtain gene-edited lines, ring-stage parasites at 5-10% parasitemia were electroporated with 50 μg of circular plasmid DNA resuspended in Cytomix. Transfected parasites were selected by culturing in the presence of 2.5 nM WR99210 (Jacobus Pharmaceuticals) for 6 days. Parasite cultures were monitored for recrudescence by microscopy and screened for successful editing by amplifying *k13* gene from recrudesced parasites using primer pair p12+p13. PCR products were sent for Sanger sequencing (*60*). Positively edited transfectants were cloned by limiting dilution to obtain clonal *k13*-edited lines before downstream phenotyping.

#### Quantification of K13 protein levels by immunoblotting

Cam3.II^R539T^, Cam3.II^R539T+A212T^ and Cam3.II^WT^ parasites were doubly synchronized using 5% D-Sorbitol to select for ring-stage parasites across 2-3 generations. For each parasite line, we harvested tightly synchronized ring and trophozoite stages on three independent occasions. Parasite pellets were collected following treatment of infected erythrocytes with 0.05% saponin, and then lysed in a mixture of SDS/Triton X-100/protease inhibitors as described previously (*63*). After centrifugation at 14,000 rpm for 10 min at 4°C to pellet cellular debris, supernatants were collected and protein concentrations determined using the DC protein assay kit (Bio-Rad). For each sample, equal amounts of protein were loaded and separated on 4–20% Tris-Glycine gels (Bio-Rad) then transferred to nitrocellulose membranes. Primary anti-K13 (mouse, monoclonal (*64*)) and anti-ERD2 (rabbit, polyclonal, MR4, BEI Resources) antibodies were used to probe K13 and ERD2 proteins. After addition of the fluorescent secondary antibodies (StarBright, Bio-Rad) bands were visualized on a ChemiDoc system (Bio-Rad). K13 and ERD2 protein levels were quantified from gel images using Image Lab 6.1 software (Bio-Rad). Total K13 protein intensity for each cell line was normalized to ERD2 loading control levels for the same line in ring and trophozoite stages separately. K13 protein levels were calculated in each line relative to those in Cam3.II^R539T^. Unpaired Welch’s t-tests were performed using Prism v8.3.1 to determine if differences in K13 protein intensity values observed between lines were statistically significant.

#### Untargeted LC-MS metabolomics of OZ439-resistant parasites

Testing for *Mycoplasma* was performed using the eMyco Detection Kit (Bocca Scientific) prior to the start of the sample collection. Mycoplasma-free parasite cultures were sorbitol synchronized across at least two generations and mid trophozoites were obtained by magnetic enrichment using MACS CS columns on the SuperMACS™ II Separator (Miltenyi Biotec, Inc.) to remove uninfected RBCs. Equal numbers of trophozoites were treated with 2 µM OZ439, 100 nM DHA, or no drug for 8 hours, then harvested for metabolites by extracting with 90% cold methanol containing the spike-in control, 0.5 µM [^13^C_4_, ^15^N_1_]-Aspartate (Cambridge Isotope) that acts as an internal LC/MS standard to correct for technical variation arising from sample processing, as described previously (*65*). For ring-stage treatments, cultures were sorbitol synchronized before treatment with 2 µM OZ439, 200 nM DHA or no drug for 8 hours and harvested as described above. Drug treatments were performed in duplicates on three independent occasions. Sample processing, quantification and analysis of metabolites including peptides was performed by LC-MS as previously described (*67*). The final processed peak intensity data of the metabolites and peptides analyzed in this study are listed in Data S1. To investigate the effect of K13 mutations on *P. falciparum* physiology, the log_2_ fold change of metabolites for the OZ439-resistant R539T+A212T double mutant vs. R539T single mutant was calculated for each experimental run using the peak areas. Statistical significance was calculated using t-tests from three independent experiments. To examine the effects of K13 mutation, we applied metabolomic set enrichment analyses and partial least squares discriminant analyses (MetaboAnalyst 3.0 package in R). For OZ439 or DHA treatments, the spectral peak areas were averaged and divided by the average of the DMSO controls for each sample type. Data were log_2_ transformed to obtain a single log_2_ fold change for each sample.

## Acknowledgments

The authors would like to acknowledge Christoph Fischli and Ursula Lehmann at Swiss TPH (Basel, Switzerland) for performing the *in vivo* antimalarial efficacy studies. We thank the Huck Institutes of Life Sciences Metabolomics Core Facility (RRID:SCR 023864) at Penn State University for maintenance of the Thermo Exactive Plus mass spectrometer. We thank Didier Leroy at the Medicines for Malaria Venture for helpful discussions.

## Author contributions

Conceptualization: SM. Methodology: CBL, SM. Formal Analysis: CBL, TY, SM. Investigation: CBL, KEW, SW, BHS, TQ, JLSS HP, SM. Visualization: CBL, SM. Supervision: ACU, ML, DAF, SM. Writing - original: CBL, SM. Writing - editing: CBL, KEW, BHS, HP, ACU, ML, DAF, SM. All authors reviewed and approved of the submitted version

## Data and materials availability

All data are available in the main text or the supplementary materials. Raw genome sequences have been deposited in NCBI Sequence Read Archive under accession numbers PRJNA1435872 and will be available upon publication of the study. Raw data for LC-MS/MS is pending submission approval at the NIH Metabolomics Workbench.

## References

1. World Health Organization, World Malaria Report 2025 (2025). https://www.who.int/teams/global-malaria-programme/reports/world-malaria-report-2025.

2. J. P. Daily, S. Parikh, Malaria. N Engl J Med 392, 1320–1333 (2025).

3. M. M. Ippolito, K. A. Moser, J.-B. B. Kabuya, C. Cunningham, J. J. Juliano, Antimalarial drug resistance and implications for the WHO global technical strategy. Curr Epidemiol Rep 8, 46–62 (2021).

4. N. J. White, K. Chotivanich, Artemisinin-resistant malaria. Clin Microbiol Rev 37, e0010924 (2024).

5. H. Noedl, Y. Se, K. Schaecher, B. L. Smith, D. Socheat, M. M. Fukuda, Evidence of artemisinin-resistant malaria in Western Cambodia. N Engl J Med 359, 2619–2620 (2008).

6. A. M. Dondorp, F. Nosten, P. Yi, D. Das, A. P. Phyo, J. Tarning, K. M. Lwin, F. Ariey, W. Hanpithakpong, S. J. Lee, P. Ringwald, K. Silamut, M. Imwong, K. Chotivanich, P. Lim, T. Herdman, S. S. An, S. Yeung, P. Singhasivanon, N. P. Day, N. Lindegardh, D. Socheat, N. J. White, Artemisinin resistance in *Plasmodium falciparum* malaria. N Engl J Med 361, 455–67 (2009).

7. M. Imwong, K. Suwannasin, S. Srisutham, R. Vongpromek, C. Promnarate, A. Saejeng, A. P. Phyo, S. Proux, T. Pongvongsa, N. Chea, O. Miotto, R. Tripura, C. Nguyen Hoang, L. Dysoley, N. Ho Dang Trung, T. J. Peto, J. J. Callery, R. W. van der Pluijm, C. Amaratunga, M. Mukaka, L. von Seidlein, M. Mayxay, N. T. Thuy-Nhien, P. N. Newton, N. P. J. Day, E. A. Ashley, F. H. Nosten, F. M. Smithuis, M. Dhorda, N. J. White, A. M. Dondorp, Evolution of multidrug resistance in *Plasmodium falciparum*: a longitudinal study of genetic resistance markers in the Greater Mekong subregion. Antimicrob Agents Chemother 65, e0112121 (2021).

8. F. Ariey, B. Witkowski, C. Amaratunga, J. Beghain, A. C. Langlois, N. Khim, S. Kim, V. Duru, C. Bouchier, L. Ma, P. Lim, R. Leang, S. Duong, S. Sreng, S. Suon, C. M. Chuor, D. M. Bout, S. Menard, W. O. Rogers, B. Genton, T. Fandeur, O. Miotto, P. Ringwald, J. Le Bras, A. Berry, J. C. Barale, R. M. Fairhurst, F. Benoit-Vical, O. Mercereau-Puijalon, D. Menard, A molecular marker of artemisinin-resistant Plasmodium falciparum malaria. Nature 505, 50–5 (2014).

9. O. Miotto, R. Amato, E. A. Ashley, B. MacInnis, J. Almagro-Garcia, C. Amaratunga, P. Lim, D. Mead, S. O. Oyola, M. Dhorda, M. Imwong, C. Woodrow, M. Manske, J. Stalker, E. Drury, S. Campino, L. Amenga-Etego, T. N. Thanh, H. T. Tran, P. Ringwald, D. Bethell, F. Nosten, A. P. Phyo, S. Pukrittayakamee, K. Chotivanich, C. M. Chuor, C. Nguon, S. Suon, S. Sreng, P. N. Newton, M. Mayxay, M. Khanthavong, B. Hongvanthong, Y. Htut, K. T. Han, M. P. Kyaw, M. A. Faiz, C. I. Fanello, M. Onyamboko, O. A. Mokuolu, C. G. Jacob, S. Takala-Harrison, C. V. Plowe, N. P. Day, A. M. Dondorp, C. C. Spencer, G. McVean, R. M. Fairhurst, N. J. White, D. P. Kwiatkowski, Genetic architecture of artemisinin-resistant *Plasmodium falciparum*. Nat Genet 47, 226–34 (2015).

10. J. Straimer, N. F. Gnadig, B. Witkowski, C. Amaratunga, V. Duru, A. P. Ramadani, M. Dacheux, N. Khim, L. Zhang, S. Lam, P. D. Gregory, F. D. Urnov, O. Mercereau-Puijalon, F. Benoit-Vical, R. M. Fairhurst, D. Menard, D. A. Fidock, K13-propeller mutations confer artemisinin resistance in Plasmodium falciparum clinical isolates. Science 347, 428–31 (2015).

11. A. J. Balmer, N. F. White, E. S. Ünlü, C. Lee, R. D. Pearson, J. Almagro-Garcia, C. Ariani, Understanding the global rise of artemisinin resistance: Insights from over 100,000 *Plasmodium falciparum* samples. eLife 14, e105544 (2025).

12. A. Uwimana, E. Legrand, B. H. Stokes, J. M. Ndikumana, M. Warsame, N. Umulisa, D. Ngamije, T. Munyaneza, J. B. Mazarati, K. Munguti, P. Campagne, A. Criscuolo, F. Ariey, M. Murindahabi, P. Ringwald, D. A. Fidock, A. Mbituyumuremyi, D. Menard, Emergence and clonal expansion of *in vitro* artemisinin-resistant *Plasmodium falciparum* kelch13 R561H mutant parasites in Rwanda. Nat Med 26, 1602–1608 (2020).

13. B. Balikagala, N. Fukuda, M. Ikeda, O. T. Katuro, S. I. Tachibana, M. Yamauchi, W. Opio, S. Emoto, D. A. Anywar, E. Kimura, N. M. Q. Palacpac, E. I. Odongo-Aginya, M. Ogwang, T. Horii, T. Mita, Evidence of artemisinin-resistant malaria in Africa. N Engl J Med 385, 1163–1171 (2021).

14. M. D. Conrad, V. Asua, S. Garg, D. Giesbrecht, K. Niaré, S. Smith, J. F. Namuganga, T. Katairo, J. Legac, R. M. Crudale, P. K. Tumwebaze, S. L. Nsobya, R. A. Cooper, M. R. Kamya, G. Dorsey, J. A. Bailey, P. J. Rosenthal, Evolution of partial resistance to artemisinins in malaria parasites in Uganda. N Engl J Med 389, 722–732 (2023).

15. P. J. Rosenthal, V. Asua, J. A. Bailey, M. D. Conrad, D. S. Ishengoma, M. R. Kamya, C. Rasmussen, F. G. Tadesse, A. Uwimana, D. A. Fidock, The emergence of artemisinin partial resistance in Africa: how do we respond? The Lancet Infectious Diseases 24, e591–e600 (2024).

16. S. R. Meshnick, Artemisinin: mechanisms of action, resistance and toxicity. International Journal for Parasitology 32, 1655–1660 (2002).

17. Z. Wang, Y. Wang, M. Cabrera, Y. Zhang, B. Gupta, Y. Wu, K. Kemirembe, Y. Hu, X. Liang, A. Brashear, S. Shrestha, X. Li, J. Miao, X. Sun, Z. Yang, L. Cui, Artemisinin resistance at the China-Myanmar border and association with mutations in the K13 propeller gene. Antimicrob Agents Chemother 59, 6952–9 (2015).

18. H. M. Ismail, V. Barton, M. Phanchana, S. Charoensutthivarakul, M. H. L. Wong, J. Hemingway, G. A. Biagini, P. M. O’Neill, S. A. Ward, Artemisinin activity-based probes identify multiple molecular targets within the asexual stage of the malaria parasites *Plasmodium falciparum* 3D7. Proc. Natl. Acad. Sci. U.S.A. 113, 2080–2085 (2016).

19. L. Tilley, J. Straimer, N. F. Gnädig, S. A. Ralph, D. A. Fidock, Artemisinin action and resistance in *Plasmodium falciparum*. Trends in Parasitology 32, 682–696 (2016).

20. J. L. Bridgford, S. C. Xie, S. A. Cobbold, C. F. A. Pasaje, S. Herrmann, T. Yang, D. L. Gillett, L. R. Dick, S. A. Ralph, C. Dogovski, N. J. Spillman, L. Tilley, Artemisinin kills malaria parasites by damaging proteins and inhibiting the proteasome. Nat Commun 9, 3801 (2018).

21. T. Yang, L. M. Yeoh, M. V. Tutor, M. W. Dixon, P. J. McMillan, S. C. Xie, J. L. Bridgford, D. L. Gillett, M. F. Duffy, S. A. Ralph, M. J. McConville, L. Tilley, S. A. Cobbold, Decreased K13 abundance reduces hemoglobin catabolism and proteotoxic stress, underpinning artemisinin resistance. Cell Rep 29, 2917–2928 e5 (2019).

22. J. Birnbaum, S. Scharf, S. Schmidt, E. Jonscher, W. A. M. Hoeijmakers, S. Flemming, C. G. Toenhake, M. Schmitt, R. Sabitzki, B. Bergmann, U. Frohlke, P. Mesen-Ramirez, A. Blancke Soares, H. Herrmann, R. Bartfai, T. Spielmann, A Kelch13-defined endocytosis pathway mediates artemisinin resistance in malaria parasites. Science 367, 51–59 (2020).

23. S. Mok, E. A. Ashley, P. E. Ferreira, L. Zhu, Z. Lin, T. Yeo, K. Chotivanich, M. Imwong, S. Pukrittayakamee, M. Dhorda, C. Nguon, P. Lim, C. Amaratunga, S. Suon, T. T. Hien, Y. Htut, M. A. Faiz, M. A. Onyamboko, M. Mayxay, P. N. Newton, R. Tripura, C. J. Woodrow, O. Miotto, D. P. Kwiatkowski, F. Nosten, N. P. J. Day, P. R. Preiser, N. J. White, A. M. Dondorp, R. M. Fairhurst, Z. Bozdech, Population transcriptomics of human malaria parasites reveals the mechanism of artemisinin resistance. Science 347, 431–435 (2015).

24. F. Rocamora, L. Zhu, K. Y. Liong, A. Dondorp, O. Miotto, S. Mok, Z. Bozdech, Oxidative stress and protein damage responses mediate artemisinin resistance in malaria parasites. PLoS Pathog 14, e1006930 (2018).

25. S. Mok, B. H. Stokes, N. F. Gnadig, L. S. Ross, T. Yeo, C. Amaratunga, E. Allman, L. Solyakov, A. R. Bottrill, J. Tripathi, R. M. Fairhurst, M. Llinas, Z. Bozdech, A. B. Tobin, D. A. Fidock, Artemisinin-resistant K13 mutations rewire *Plasmodium falciparum’s* intra-erythrocytic metabolic program to enhance survival. Nat Commun 12, 530 (2021).

26. M. R. Rosenthal, S. Vijayrajratnam, T. M. Firestone, C. L. Ng, Enhanced cell stress response and protein degradation capacity underlie artemisinin resistance in *Plasmodium falciparum*. mSphere 9, e00371–24 (2024).

27. J. L. Vennerstrom, S. Arbe-Barnes, R. Brun, S. A. Charman, F. C. K. Chiu, J. Chollet, Y. Dong, A. Dorn, D. Hunziker, H. Matile, K. McIntosh, M. Padmanilayam, J. Santo Tomas, C. Scheurer, B. Scorneaux, Y. Tang, H. Urwyler, S. Wittlin, W. N. Charman, Identification of an antimalarial synthetic trioxolane drug development candidate. Nature 430, 900–904 (2004).

28. C. S. Perry, S. A. Charman, R. J. Prankerd, F. C. K. Chiu, Y. Dong, J. L. Vennerstrom, W. N. Charman, Chemical kinetics and aqueous degradation pathways of a new class of synthetic ozonide antimalarials. Journal of Pharmaceutical Sciences 95, 737–747 (2006).

29. M. Kaiser, S. Wittlin, A. Nehrbass-Stuedli, Y. Dong, X. Wang, A. Hemphill, H. Matile, R. Brun, J. L. Vennerstrom, Peroxide bond-dependent antiplasmodial specificity of artemisinin and OZ277 (RBx11160). Antimicrob Agents Chemother 51, 2991–2993 (2007).

30. J. Jourdan, H. Matile, E. Reift, O. Biehlmaier, Y. Dong, X. Wang, P. Mäser, J. L. Vennerstrom, S. Wittlin, Monoclonal antibodies that recognize the alkylation signature of antimalarial ozonides OZ277 (Arterolane) and OZ439 (Artefenomel). ACS Infect. Dis. 2, 54–61 (2016).

31. C. Wei, C.-X. Zhao, S. Liu, J.-H. Zhao, Z. Ye, H. Wang, S.-S. Yu, C.-J. Zhang, Activity-based protein profiling reveals that secondary-carbon-centered radicals of synthetic 1,2,4-trioxolanes are predominately responsible for modification of protein targets in malaria parasites. Chem. Commun. 55, 9535–9538 (2019).

32. M. Nguyen, L. Paloque, J. Manaranche, M. Chabbert, A. Hamouy, M. Laurent, J.-M. Augereau, C. Claparols, A. Robert, F. Benoit-Vical, Reductive activation of artefenomel (OZ439) by Fe(II)-heme, related to its antimalarial activity. ACS Infect. Dis. 11, 216–225 (2025).

33. C. Giannangelo, F. J. I. Fowkes, J. A. Simpson, S. A. Charman, D. J. Creek, Ozonide antimalarial activity in the context of artemisinin-resistant malaria. Trends Parasitol 35, 529–543 (2019).

34. S. A. Charman, S. Arbe-Barnes, I. C. Bathurst, R. Brun, M. Campbell, W. N. Charman, F. C. Chiu, J. Chollet, J. C. Craft, D. J. Creek, Y. Dong, H. Matile, M. Maurer, J. Morizzi, T. Nguyen, P. Papastogiannidis, C. Scheurer, D. M. Shackleford, K. Sriraghavan, L. Stingelin, Y. Tang, H. Urwyler, X. Wang, K. L. White, S. Wittlin, L. Zhou, J. L. Vennerstrom, Synthetic ozonide drug candidate OZ439 offers new hope for a single-dose cure of uncomplicated malaria. Proc Natl Acad Sci U S A 108, 4400–5 (2011).

35. Y. Dong, X. Wang, S. Kamaraj, V. J. Bulbule, F. C. K. Chiu, J. Chollet, M. Dhanasekaran, C. D. Hein, P. Papastogiannidis, J. Morizzi, D. M. Shackleford, H. Barker, E. Ryan, C. Scheurer, Y. Tang, Q. Zhao, L. Zhou, K. L. White, H. Urwyler, W. N. Charman, H. Matile, S. Wittlin, S. A. Charman, J. L. Vennerstrom, Structure–activity relationship of the antimalarial ozonide artefenomel (OZ439). J. Med. Chem. 60, 2654–2668 (2017).

36. N. Valecha, S. Looareesuwan, A. Martensson, S. M. Abdulla, S. Krudsood, N. Tangpukdee, S. Mohanty, S. K. Mishra, P. K. Tyagi, S. K. Sharma, J. Moehrle, A. Gautam, A. Roy, J. K. Paliwal, M. Kothari, N. Saha, A. P. Dash, A. Björkman, Arterolane, a new synthetic trioxolane for treatment of uncomplicated *Plasmodium falciparum* malaria: A phase II, multicenter, randomized, dose-finding clinical trial. CLIN INFECT DIS 51, 684–691 (2010).

37. N. Saha, J. J. Moehrle, A. Zutshi, P. Sharma, P. Kaur, S. S. Iyer, Safety, tolerability and pharmacokinetic profile of single and multiple oral doses of arterolane (RBx11160) maleate in healthy subjects. The Journal of Clinical Pharmacology 54, 386–393 (2014).

38. O. A. Toure, S. Rulisa, A. R. Anvikar, B. S. Rao, P. Mishra, R. K. Jalali, S. Arora, A. Roy, N. Saha, S. S. Iyer, P. Sharma, N. Valecha, Efficacy and safety of fixed dose combination of arterolane maleate and piperaquine phosphate dispersible tablets in paediatric patients with acute uncomplicated *Plasmodium falciparum* malaria: a phase II, multicentric, open-label study. Malar J 14, 469 (2015).

39. J. J. Moehrle, S. Duparc, C. Siethoff, P. L. M. Van Giersbergen, J. C. Craft, S. Arbe-Barnes, S. A. Charman, M. Gutierrez, S. Wittlin, J. L. Vennerstrom, First-in-man safety and pharmacokinetics of synthetic ozonide OZ439 demonstrates an improved exposure profile relative to other peroxide antimalarials. Brit J Clinical Pharma 75, 535–548 (2013).

40. A. P. Phyo, P. Jittamala, F. H. Nosten, S. Pukrittayakamee, M. Imwong, N. J. White, S. Duparc, F. Macintyre, M. Baker, J. J. Mohrle, Antimalarial activity of artefenomel (OZ439), a novel synthetic antimalarial endoperoxide, in patients with *Plasmodium vivax* malaria: an open-label phase 2 trial. Lancet Infect Dis 16, 61–69 (2016).

41. F. Baumgärtner, J. Jourdan, C. Scheurer, B. Blasco, B. Campo, P. Mäser, S. Wittlin, *In vitro* activity of anti-malarial ozonides against an artemisinin-resistant isolate. Malar J 16, 45 (2017).

42. J. Straimer, N. F. Gnadig, B. H. Stokes, M. Ehrenberger, A. A. Crane, D. A. Fidock, Plasmodium falciparum K13 mutations differentially impact ozonide susceptibility and parasite fitness in vitro. mBio 8 (2017).

43. A. Walz, D. Leroy, N. Andenmatten, P. Maser, S. Wittlin, Anti-malarial ozonides OZ439 and OZ609 tested at clinically relevant compound exposure parameters in a novel ring-stage survival assay. Malar J 18, 427 (2019).

44. F. Macintyre, Y. Adoke, A. B. Tiono, T. T. Duong, G. Mombo-Ngoma, M. Bouyou-Akotet, H. Tinto, Q. Bassat, S. Issifou, M. Adamy, H. Demarest, S. Duparc, D. Leroy, B. E. Laurijssens, S. Biguenet, A. Kibuuka, A. K. Tshefu, M. Smith, C. Foster, I. Leipoldt, P. G. Kremsner, B. Q. Phuc, A. Ouedraogo, M. Ramharter, O. Z.-P. S. Group, A randomised, double-blind clinical phase II trial of the efficacy, safety, tolerability and pharmacokinetics of a single dose combination treatment with artefenomel and piperaquine in adults and children with uncomplicated *Plasmodium falciparum* malaria. BMC Med 15, 181 (2017).

45. Y. Adoke, R. Zoleko-Manego, S. Ouoba, A. B. Tiono, G. Kaguthi, J. E. Bonzela, T. T. Duong, A. Nahum, M. Bouyou-Akotet, B. Ogutu, A. Ouedraogo, F. Macintyre, A. Jessel, B. Laurijssens, M. H. Cherkaoui-Rbati, C. Cantalloube, A. C. Marrast, R. Bejuit, D. White, T. N. C. Wells, F. Wartha, D. Leroy, A. Kibuuka, G. Mombo-Ngoma, D. Ouattara, I. Mugenya, B. Q. Phuc, F. Bohissou, D. P. Mawili-Mboumba, F. Olewe, I. Soulama, H. Tinto, the FALCI Study Group, M. Ramharter, D. Nahum, H. Zohou, I. Nzwili, J. M. Ongecha, R. Thompson, J. Kiwalabye, A. Diarra, A. S. Coulibaly, E. C. Bougouma, D. G. Kargougou, M. Tegneri, C. Castin Vuillerme, E. Djeriou, A. F. Ansary, A randomized, double-blind, phase 2b study to investigate the efficacy, safety, tolerability and pharmacokinetics of a single-dose regimen of ferroquine with artefenomel in adults and children with uncomplicated *Plasmodium falciparum* malaria. Malar J 20, 222 (2021).

46. A. Gansané, L. F. Moriarty, D. Ménard, I. Yerbanga, E. Ouedraogo, P. Sondo, R. Kinda, C. Tarama, E. Soulama, M. Tapsoba, D. Kangoye, C. S. Compaore, O. Badolo, B. Dao, S. Tchwenko, H. Tinto, I. Valea, Anti-malarial efficacy and resistance monitoring of artemether-lumefantrine and dihydroartemisinin-piperaquine shows inadequate efficacy in children in Burkina Faso, 2017–2018. Malar J 20, 48 (2021).

47. M. T. Klope, J. A. Tapia Cardona, J. Chen, R. L. Gonciarz, K. Cheng, P. Jaishankar, J. Kim, J. Legac, P. J. Rosenthal, A. R. Renslo, Synthesis and *in vivo* profiling of desymmetrized antimalarial trioxolanes with diverse carbamate side chains. ACS Med. Chem. Lett. 15, 1764–1770 (2024).

48. M. T. Klope, P. Talukder, B. R. Blank, S. Chelebieva, J. Chen, S. D. Fontaine, R. L. Gonciarz, P. Jaishankar, G. J. Lee, J. Legac, V. Mathur, A. Narayan, M. Okitwi, S. Orena, N. S. Settineri, J. A. Tapia, Y. Taremwa, P. K. Tumwebaze, A. Vinod, J. N. Burrows, P. J. Rosenthal, R. A. Cooper, A. R. Renslo, Identifying a next-generation antimalarial trioxolane in a landscape of artemisinin partial resistance. Sci. Adv. 11, eads9168 (2025).

49. V. Wasakul, A. Disratthakit, M. Mayxay, K. Chindavongsa, V. Sengsavath, N. Thuy-Nhien, R. D. Pearson, S. Phalivong, S. Xayvanghang, R. J. Maude, S. Goncalves, N. P. Day, P. N. Newton, E. A. Ashley, D. P. Kwiatkowski, A. M. Dondorp, O. Miotto, Malaria outbreak in Laos driven by a selective sweep for *Plasmodium falciparum* kelch13 R539T mutants: a genetic epidemiology analysis. Lancet Infect Dis, doi: 10.1016/S1473-3099(22)00697-1 (2022).

50. J. S. McCarthy, M. Baker, P. O’Rourke, L. Marquart, P. Griffin, R. Hooft van Huijsduijnen, J. J. Mohrle, Efficacy of OZ439 (artefenomel) against early *Plasmodium falciparum* blood-stage malaria infection in healthy volunteers. J Antimicrob Chemother 71, 2620–7 (2016).

51. Malaria Genomic Epidemiology Network (MalariaGEN), M. M. Abdel Hamid, M. H. Abdelraheem, D. O. Acheampong, I. Adam, P. Aide, O. Ajibaye, M. Ali, J. Almagro-Garcia, A. Amambua-Ngwa, L. Amenga-Etego, I. Aniebo, E. Aninagyei, F. Ansah, T. O. Apinjoh, C. V. Ariani, S. Auburn, G. A. Awandare, A. Balmer, P. Bejon, S. Boene, G. Bwire, B. Candrinho, A. Chidimatembue, K. Chindavongsa, K. Comiche, D. Conway, A. Dara, M. Diakite, A. Djimde, A. Dondorp, S. Doumbia, E. Drury, C. A. Fanello, M. Ferdig, K. Figueroa, D. Gamboa, L. Golassa, S. Gonçalves, M. D. A. Guindo, M. Hamaluba, B. Hanboonkunupakarn, K. Howe, M. Hussien, M. Imwong, D. Ishengoma, J. Jeans, A. Kabaghe, A. Kamuhabwa, J.-M. Kindermans, D. S. Konate, D. P. Kwiatkowski, C. Lee, S. K. Lee, S. J. Lee, B. Ley, A. Llanos-Cuentas, J. Marfurt, G. Matambisso, R. R. Maude, R. J. Maude, A. Mayor, M. Mayxay, O. Maïga-Ascofaré, R. S. McCann, A. Miles, O. Miotto, A. O. Mohamed, C. M. Morang’a, K. Murie, B. E. Ngasala, T.-N. Nguyen, O. Nolasco, F. Nosten, R. Noviyanti, Í. O’Connor, M. Oboh, L. I. Ochola-Oyier, C. Olufunke Falade, A. Olukosi, A. Olumide, F. I. Olusola, M. A. Onyamboko, E. C. Oriero, W. A. Oyibo, D. Pannebaker, R. D. Pearson, K. Phiri, R. W. Van Der Pluijm, R. N. Price, H. H. Quang, V. Rajkumar Devaraju, M. Randrianarivelojosia, L. Ranford-Cartwright, J. C. Rayner, E. Rovira-Vallbona, K. Rowlands, V. Ruano-Rubio, J. F. Sanchez, F. Saúte, S. Shettima, C. Da Silva, V. J. Simpson, S. Suddaby, W. Takken, A. M. Thu, M. Toure, E. Unlu, H. O. Valdivia, M. Van Vugt, N. Waithira, T. Wellems, J. Wendler, N. White, R. Wuendrich Ogidan, Pf8: an open dataset of Plasmodium falciparum genome variation in 33,325 worldwide samples. Wellcome Open Res 10, 325 (2025).

52. B. Witkowski, J. Lelièvre, M. J. López Barragán, V. Laurent, X. Su, A. Berry, F. Benoit-Vical, Increased tolerance to artemisinin in *Plasmodium falciparum* Is mediated by a quiescence mechanism. Antimicrob Agents Chemother 54, 1872–1877 (2010).

53. D. J. Creek, W. N. Charman, F. C. K. Chiu, R. J. Prankerd, Y. Dong, J. L. Vennerstrom, S. A. Charman, Relationship between antimalarial activity and heme alkylation for spiro- and dispiro-1,2,4-trioxolane antimalarials. Antimicrob Agents Chemother 52, 1291–1296 (2008).

54. C. Giannangelo, D. Anderson, X. Wang, J. L. Vennerstrom, S. A. Charman, D. J. Creek, Ozonide antimalarials alkylate heme in the malaria parasite *Plasmodium falciparum*. ACS Infect. Dis. 5, 2076–2086 (2019).

55. C. Giannangelo, G. Siddiqui, A. De Paoli, B. M. Anderson, L. E. Edgington-Mitchell, S. A. Charman, D. J. Creek, System-wide biochemical analysis reveals ozonide antimalarials initially act by disrupting *Plasmodium falciparum* haemoglobin digestion. PLoS Pathog 16, e1008485 (2020).

56. S. Muller, Role and Regulation of glutathione metabolism in *Plasmodium falciparum*. Molecules 20, 10511–34 (2015).

57. J. Tripathi, M. Stoklasa, S. Nayak, K. En Low, E. Qian Hui Lee, Q. H. Duong Tien, L. Rénia, B. Malleret, Z. Bozdech, Author Correction: The artemisinin-induced dormant stages of *Plasmodium falciparum* exhibit hallmarks of cellular quiescence/senescence and drug resilience. Nat Commun 15, 8460 (2024).

58. I. Angulo-Barturen, M. B. Jimenez-Diaz, T. Mulet, J. Rullas, E. Herreros, S. Ferrer, E. Jimenez, A. Mendoza, J. Regadera, P. J. Rosenthal, I. Bathurst, D. L. Pompliano, F. Gomez de las Heras, D. Gargallo-Viola, A murine model of falciparum-malaria by *in vivo* selection of competent strains in non-myelodepleted mice engrafted with human erythrocytes. PLoS One 3, e2252 (2008).

59. M. B. Jimenez-Diaz, T. Mulet, S. Viera, V. Gomez, H. Garuti, J. Ibanez, A. Alvarez-Doval, L. D. Shultz, A. Martinez, D. Gargallo-Viola, I. Angulo-Barturen, Improved murine model of malaria using *Plasmodium falciparum* competent strains and non-myelodepleted NOD-scid IL2Rgammanull mice engrafted with human erythrocytes. Antimicrob Agents Chemother 53, 4533–6 (2009).

60. M. Kanai, T. Yeo, V. Asua, P. J. Rosenthal, D. A. Fidock, S. Mok, Comparative analysis of *Plasmodium falciparum* genotyping via SNP detection, microsatellite profiling, and whole-genome sequencing. Antimicrob Agents Chemother 66, e0116321 (2022).

61. B. Witkowski, C. Amaratunga, N. Khim, S. Sreng, P. Chim, S. Kim, P. Lim, S. Mao, C. Sopha, B. Sam, J. M. Anderson, S. Duong, C. M. Chuor, W. R. Taylor, S. Suon, O. Mercereau-Puijalon, R. M. Fairhurst, D. Menard, Novel phenotypic assays for the detection of artemisinin-resistant *Plasmodium falciparum* malaria in Cambodia: in-vitro and ex-vivo drug-response studies. Lancet Infect Dis 13, 1043–9 (2013).

62. C. Amaratunga, A. T. Neal, R. M. Fairhurst, Flow cytometry-based analysis of artemisinin-resistant *Plasmodium falciparum* in the ring-stage survival assay. Antimicrob Agents Chemother 58, 4938–40 (2014).

63. B. H. Stokes, S. K. Dhingra, K. Rubiano, S. Mok, J. Straimer, N. F. Gnädig, I. Deni, K. A. Schindler, J. R. Bath, K. E. Ward, J. Striepen, T. Yeo, L. S. Ross, E. Legrand, F. Ariey, C. H. Cunningham, I. M. Souleymane, A. Gansané, R. Nzoumbou-Boko, C. Ndayikunda, A. M. Kabanywanyi, A. Uwimana, S. J. Smith, O. Kolley, M. Ndounga, M. Warsame, R. Leang, F. Nosten, T. J. Anderson, P. J. Rosenthal, D. Ménard, D. A. Fidock, *Plasmodium falciparum* K13 mutations in Africa and Asia impact artemisinin resistance and parasite fitness. eLife 10, e66277 (2021).

64. N. F. Gnädig, B. H. Stokes, R. L. Edwards, G. F. Kalantarov, K. C. Heimsch, M. Kuderjavy, A. Crane, M. C. S. Lee, J. Straimer, K. Becker, I. N. Trakht, A. R. Odom John, S. Mok, D. A. Fidock, Insights into the intracellular localization, protein associations and artemisinin resistance properties of *Plasmodium falciparum* K13. PLoS Pathog 16, e1008482 (2020).

65. E. L. Allman, H. J. Painter, J. Samra, M. Carrasquilla, M. Llinás, Metabolomic profiling of the Malaria Box Reveals Antimalarial Target Pathways. Antimicrob Agents Chemother 60, 6635–6649 (2016).

66. E. L. Allman, H. J. Painter, J. Samra, M. Carrasquilla, M. Llinás, Metabolomic Profiling of the Malaria Box Reveals Antimalarial Target Pathways. *Antimicrobial Agents and Chemotherapy*, AAC.01224–16 (2016).

67. J. Okombo, S. Mok, T. Qahash, T. Yeo, J. Bath, L. M. Orchard, E. Owens, I. Koo, I. Albert, M. Llinas, D. A. Fidock, Piperaquine-resistant PfCRT mutations differentially impact drug transport, hemoglobin catabolism and parasite physiology in *Plasmodium falciparum* asexual blood stages. PLoS Pathog 18, e1010926 (2022).

