## Supplementary Figures 1-8 and Table S1 for "Double Mutations in *Plasmodium falciparum* Kelch13 drive resistance to next-generation artemisinin derivatives in malaria parasites"

Christopher Bower-Lepts *et al.*

**This PDF file includes:**

Figs. S1 to S8
Tables S1

**Other Supplementary Materials for this manuscript include the following:**
Data S1

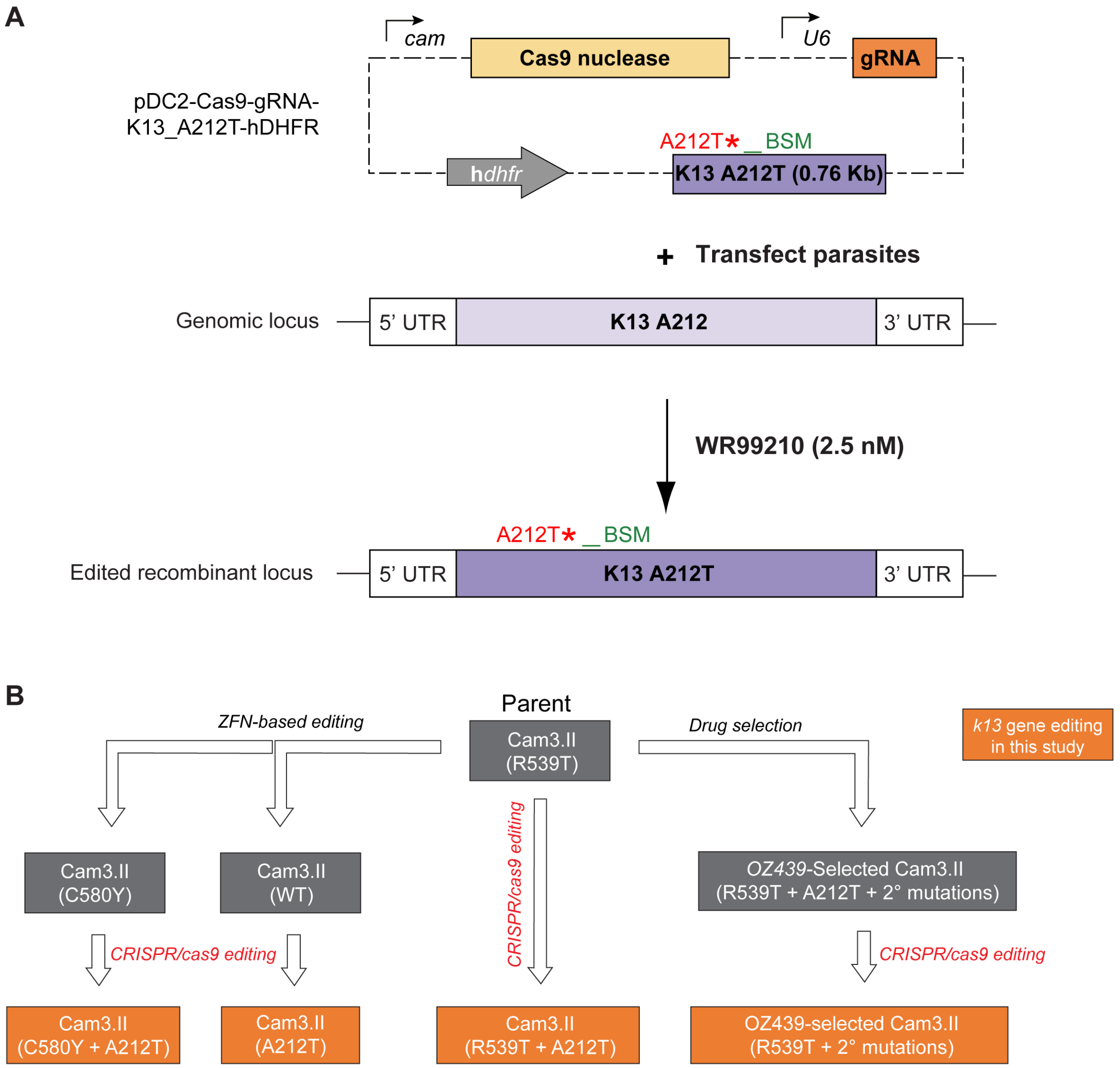


**Fig. S1. Gene-editing approach to assess the impact of *k13* mutations on survival to OZ439 in Cam3.II parasites. (A)** Schematic representation of the CRISPR/cas9 approach utilized to generate *k13*-edited *P. falciparum* parasites in this study. The pDC2-cam-coSpCas9-U6-gRNA-h*dhfr* all in one plasmid was used to introduce or remove the A212T locus in Cam3.II parasite clones. **(B)** Overview of the *k13* mutant lines subjected to phenotyping assays and the parental, gene-edited or drug-selected parasites from which they are derived. *k13*-edited lines generated in this study using CRISPR/cas9 are highlighted in orange. For the list of gene-edited parasite lines generated in this study refer to Table 1. BSM: Silent binding site mutations.

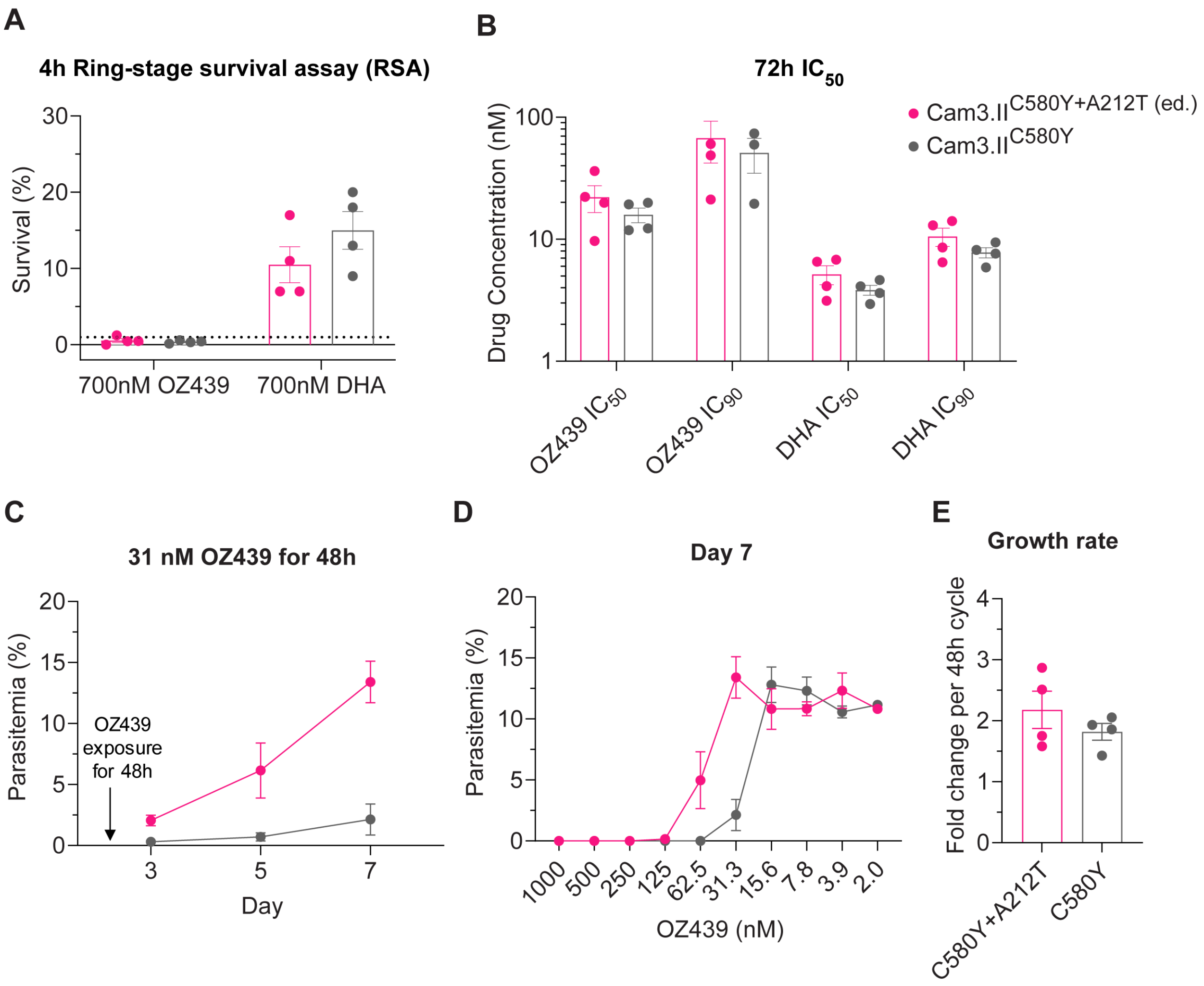


**Fig. S2. K13 A212T conferred OZ439 resistance on the Cam3.II K13 C580Y background.** **(A)** Percentage ring-stage survival of Cam3.II$^{C580Y+A212T}$ and Cam3.II$^{C580Y}$ following 4-hour exposure to 700 nM OZ439 or DHA. The dashed line 1% survival denotes the threshold for *in vitro* ART resistance in ring-stage survival assays. **(B)** $IC_{50}$ and $IC_{90}$ values of OZ439 and DHA determined from 72-hour dose-response assays using Cam3.II$^{C580Y+A212T}$ and Cam3.II$^{C580Y}$ parasites. **(C)** Parasitemia of Cam3.II$^{C580Y+A212T}$ and Cam3.II$^{C580Y}$ parasite lines measured at 48-hour intervals from 3-7 days following 48-hour treatment with 31 nM OZ439. **(D)** Parasitemia of Cam3.II$^{C580Y+A212T}$ and Cam3.II$^{C580Y}$ measured on day 7 following 48-hour treatment across a range of OZ439 concentrations. **(E)** Growth rate of Cam3.II$^{C580Y+A212T}$ and Cam3.II$^{C580Y}$ expressed as fold-change per 48-hour cycle. Points in each graph represent mean values determined across three independent experiments with technical duplicates. Error bars in each graph represent the SEM.

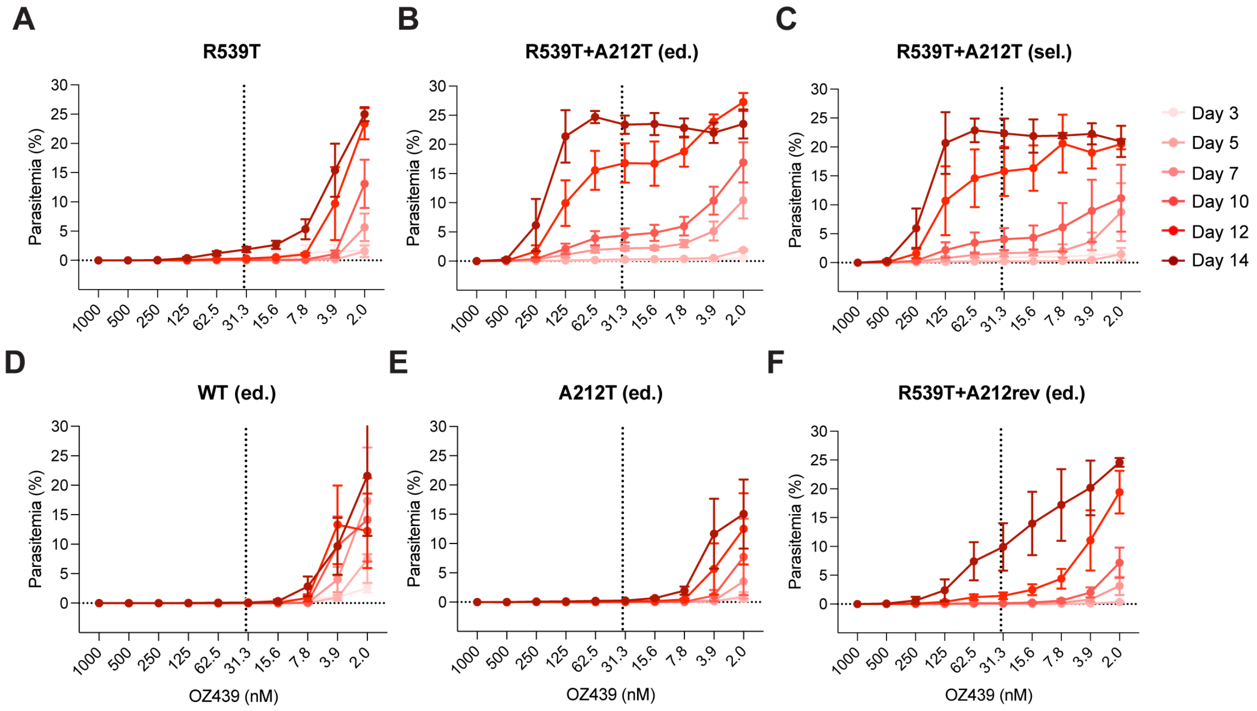


**Fig. S3. The K13 A212T+R539T parasites demonstrated accelerated recovery following 48-hour exposure to OZ439.** (**A** to **F**) Parasitemia of resistance-selected, *k13*-edited and wild-type Cam3.II parasites following exposure to a range of OZ439 concentrations for 48 hours measured at day 3-14 following drug exposure. Individual graphs represent different test lines including (A) R539T, (B) R539T+A212T (ed.), (C) R539T+A212T (sel.), (D) Wild-type, (E) A212T (ed.) and (F) R539T+A212Trev (ed.). Lines are coloured according to the day at which parasitemia was recorded. The vertical dashed line in each graph denotes the clinically relevant OZ439 concentration of 34 nM. Points in each graph represent mean values determined across three independent experiments with technical duplicates. Error bars in each graph represent SEM.

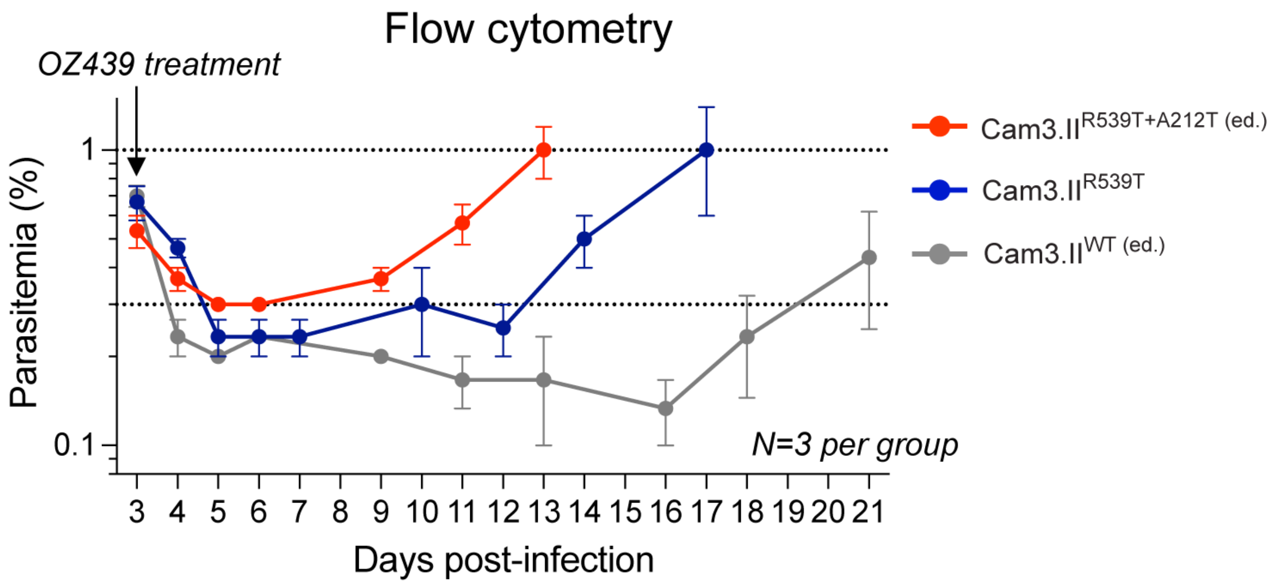


**Fig. S4. *In vivo* survival of *k13*-edited Cam3.II lines to 10 mg/kg OZ439 administered in NOD scid gamma mice determined by flow cytometry.** Percentage parasitemia of Cam3.II$^{R539T+A212T}$, Cam3.II$^{R539T}$ and Cam3.II$^{WT}$ parasite lines measured for up to 21 days following administration of 10 mg/kg OZ439 mesylate. Values represent the mean parasitemia in the peripheral blood of three individual mice obtained from flow cytometric analysis of 10,000 erythrocytes. Error bars represent the SEM.

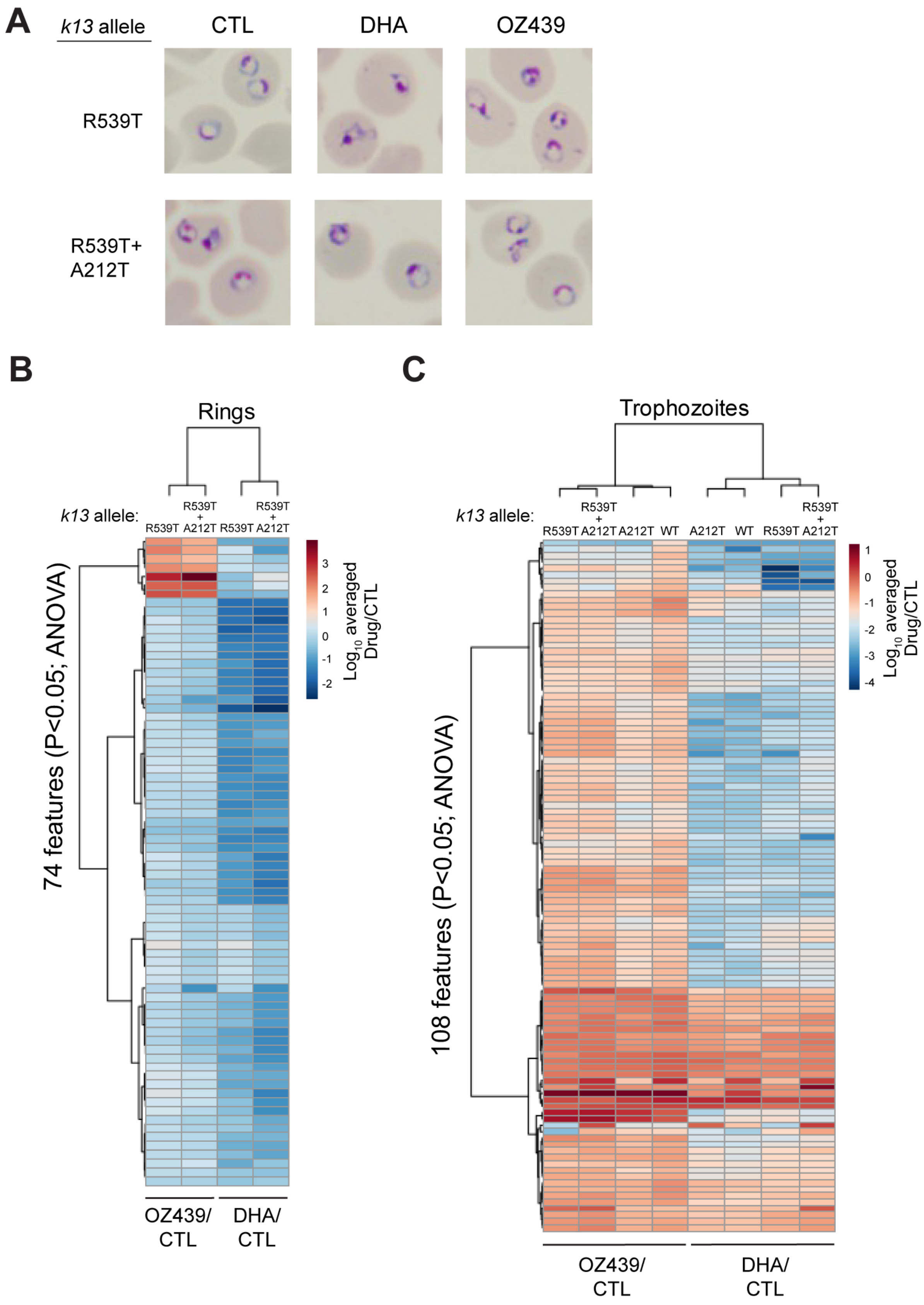
A
k13 allele
CTL
DHA
OZ439
R539T
R539T+
A212T
B
Rings
k13 allele:
R539T
R539T + A212T
R539T
R539T + A212T
74 features (P<0.05; ANOVA)
Log10 averaged Drug/CTL
3
2
1
0
-1
-2
OZ439/ CTL
DHA/ CTL
C
Trophozoites
k13 allele:
R539T
R539T + A212T
A212T
WT
A212T
WT
R539T
R539T + A212T
108 features (P<0.05; ANOVA)
Log10 averaged Drug/CTL
1
0
-1
-2
-3
-4
OZ439/ CTL
DHA/ CTL

**Fig. S5. Treatment with OZ439 and DHA causes peptide downregulation across all K13-edited lines in ring or trophozoite stages. (A)** Giemsa-stained parasite blood smears of Cam3.II$^{R539T}$ and Cam3.II$^{R539T+A212T}$ parasite lines following treatment of ring stages with 2 µM OZ439, 100 nM DHA, or no drug for 8h. (**B**) Heat map depicts the significantly differentially expressed peptides between the Cam3.II$^{R539T+A212T}$ vs Cam3.II$^{R539T}$ lines at the ring stage upon DHA or OZ439 treatment (determined using ANOVA ($P<0.05$)). Values represent the drug-treated divided by untreated control samples. Red and blue colors represent the up- or downregulated peptides respectively. (**C**) Heat map depicts the significantly differentially expressed peptides between the Cam3.II$^{R539T+A212T}$, Cam3.II$^{A212T}$ and Cam3.II$^{WT}$ vs Cam3.II$^{R539T}$ lines at the trophozoite stage upon DHA or OZ439 treatment (determined using ANOVA ($P<0.05$)). Values represent the drug-treated divided by untreated control samples. Red and blue colors represent the up- or downregulated peptides respectively.

### Pathway analysis of DE metabolites (non-peptides) in Cam3.II$^{R539T}$ and Cam3.II$^{R539T+A212T}$ in ring stages

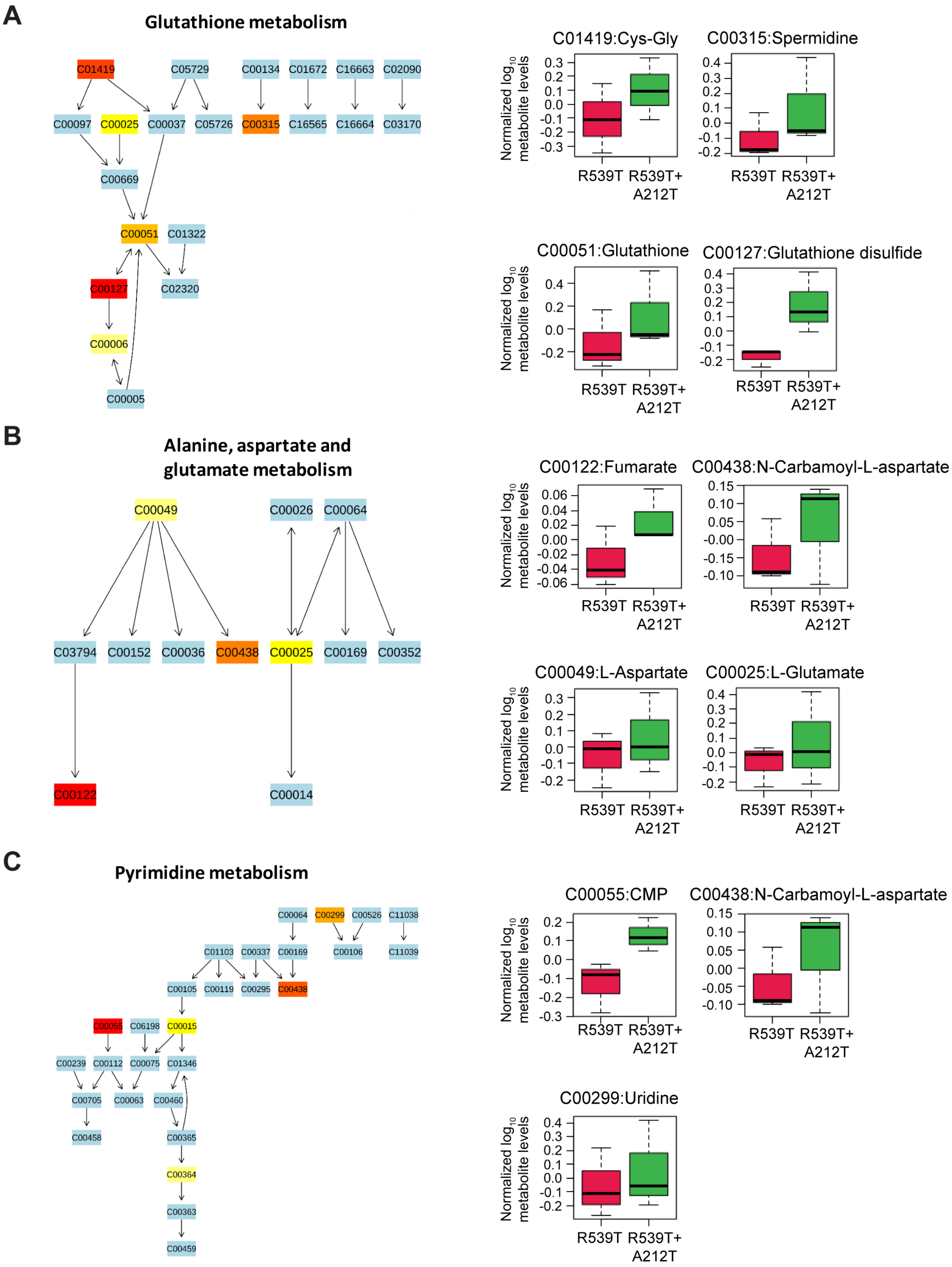

**Fig. S6. Metabolic pathways and the corresponding metabolites significantly enriched in Cam3.II$^{R539T+A212T}$ relative to Cam3.II$^{R539T}$ in ring stage. (A** to **C)** Schematic of the metabolites belonging to the glutathione (A), alanine, aspartate and glutamate (B) and pyrimidine (C) pathways. In schematics, orange and red represents differential abundance of individual metabolites between the lines, yellow indicates no significant difference. Box and whisker graphs depict normalized $log_2$-transformed levels of individual metabolites within each pathway in the Cam3.II$^{R539T}$ and Cam3.II$^{R539T+A212T}$ parasite lines.

### Pathway analysis of DE metabolites (non-peptides) in Cam3.II$^{R539T}$ and Cam3.II$^{R539T+A212T}$ in trophozoite stages

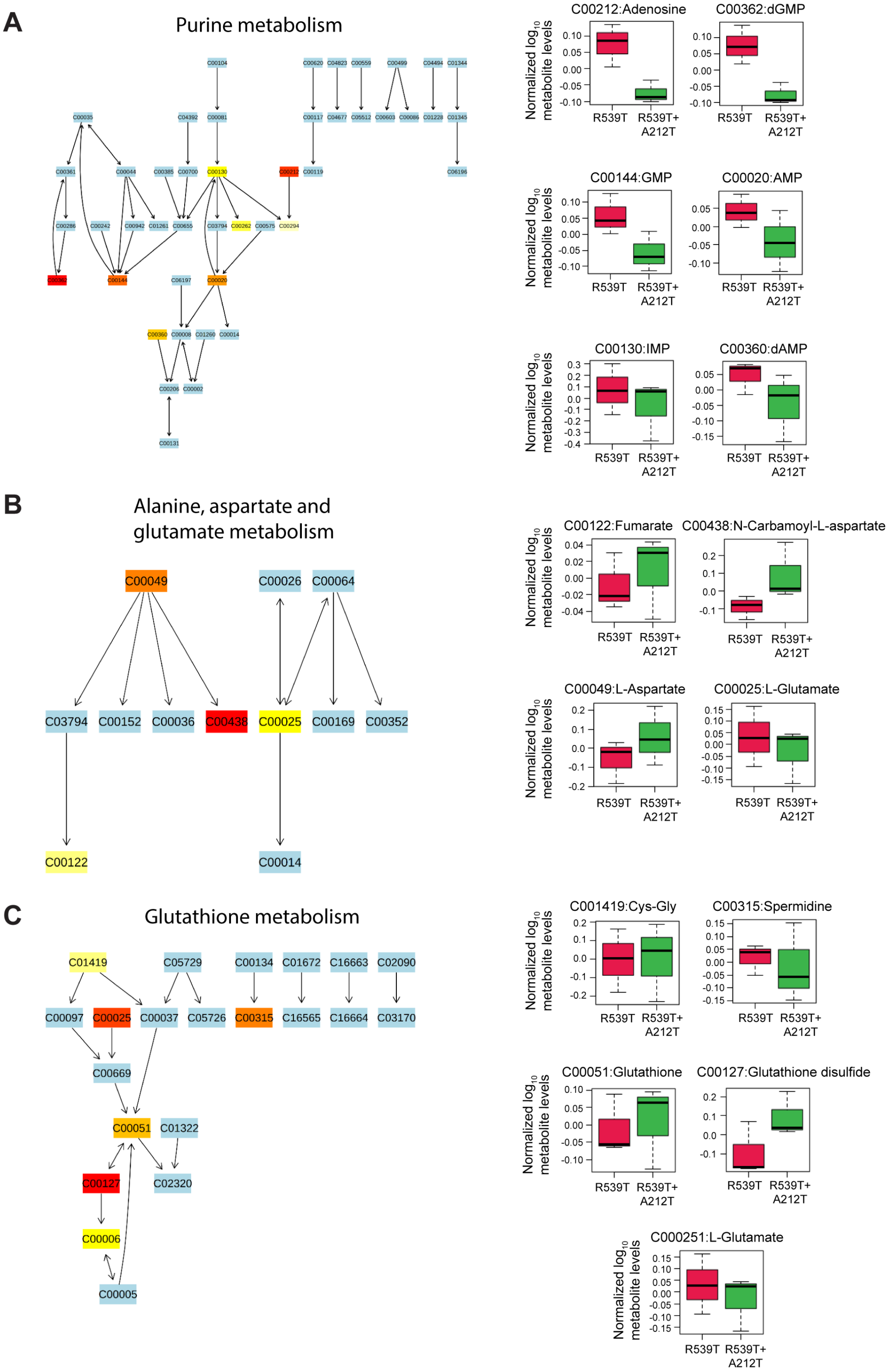

**Fig. S7. Metabolic pathways and the corresponding metabolites significantly enriched in Cam3.II$^{R539T+A212T}$ relative to Cam3.II$^{R539T}$ in trophozoite stage. (A** to **C)** Schematic of the metabolites belonging to the purine (A), alanine, aspartate and glutamate (B) and glutathione (C) pathways. In schematics, orange and red represents differential abundance of individual metabolites between the lines, yellow indicates no significant difference. Box and whisker graphs depict normalized $log_2$-transformed levels of individual metabolites within each pathway in the Cam3.II$^{R539T}$ and Cam3.II$^{R539T+A212T}$ parasite lines.

**Pathway analysis of DE metabolites (non-peptides) in Cam3.II$^{A212T}$ and Cam3.II$^{WT}$ in trophozoite stages**

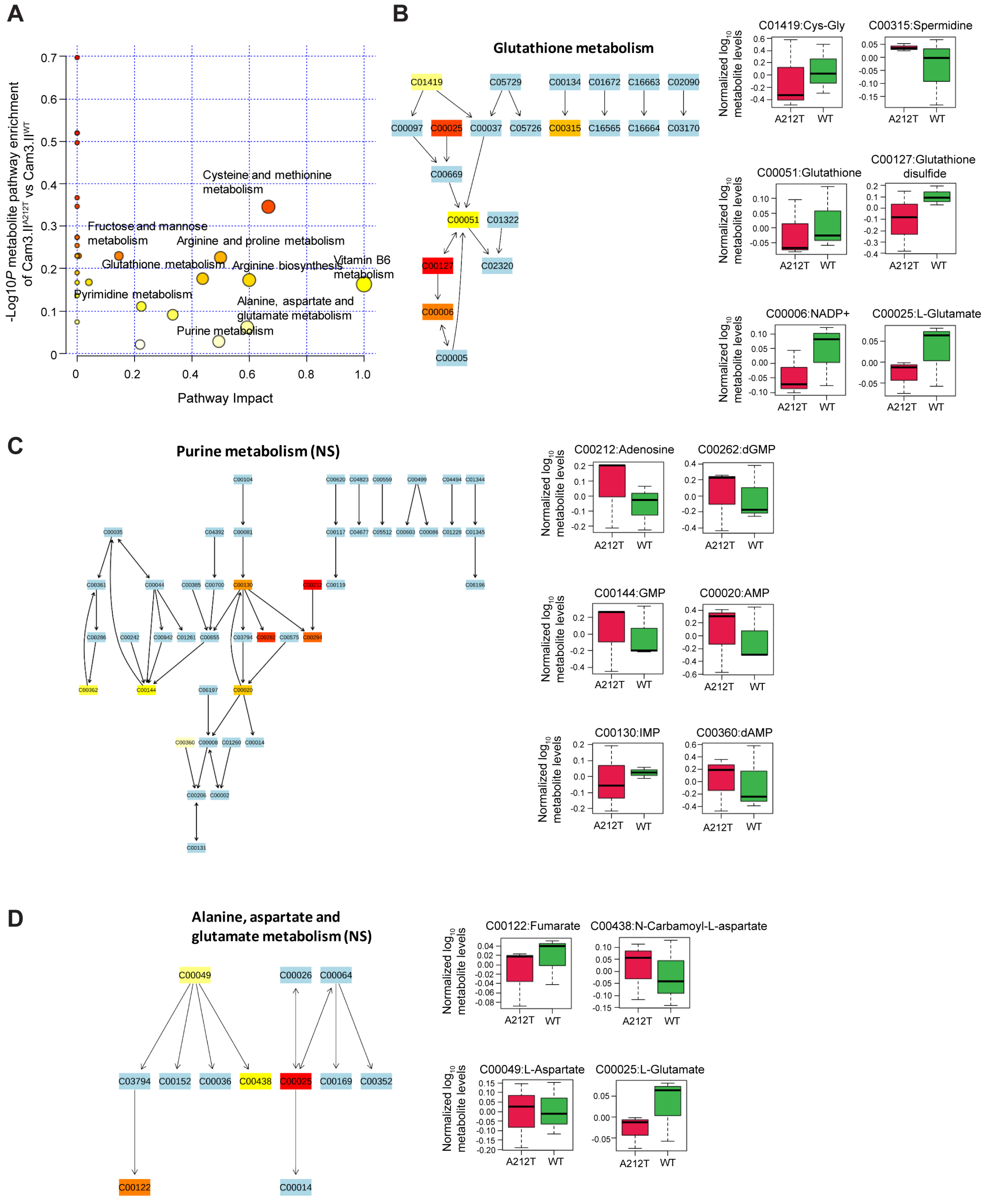

**Fig. S8. Metabolic pathways and the corresponding metabolites significantly enriched in Cam3.II$^{A212T}$ relative to Cam3.II$^{WT}$ in trophozoite stage. (A)** Metabolic pathway enrichment analysis in Cam3.II$^{A212T}$ parasites relative to Cam3.II$^{WT}$ parasites determined in the absence of drug treatment. Circle size represents pathway impact value and are colored according to the enrichment $-\log_{10} P$ score. **(B** to **D)** Schematic of the metabolites belonging to the pathways which were significantly enriched in Cam3.II$^{R539T+A212T}$ double mutants include glutathione (B), purine (C) and alanine, aspartate and glutamate (D) pathways. Metabolites in these pathways were not elevated in Cam3.II$^{A212T}$ parasites indicating that their enrichment was specific to OZ439-resistant double mutants (refer to Fig. 6). In schematics, orange and red represents differential abundance of individual metabolites between the lines, yellow indicates no significant difference. Box and whisker graphs depict normalized $\log_2$-transformed levels of individual metabolites within each pathway in the Cam3.II$^{A212T}$ and Cam3.II$^{WT}$ parasite lines.

**Table S1. List of oligonucleotides for *k13* gene-editing used in this study.**

| ID | Nucleotide sequence (5'-3') | Description* | Lab ID |
|---|---|---|---|
| **p1** | TATTTAAGAATTACATTTATTAAT | *k13* gRNA fwd | p7927 |
| **p2** | AAACATTAATAAATGTAATTCTTA | *k13* gRNA rev | p7928 |
| **p3** | GTAGCGAGAATGATTCTAATTC | *k13* CRISPR/Cas9 donor amplification fwd | p7940 |
| **p4** | TGTTTATAACCATTAGATATATCAATATC | *k13* CRISPR/Cas9 donor amplification rev | p7941 |
| **p5** | GAAAATATGGTAGGTGATTTAAGAATTACATTcATcAAcTGGTTAAAAAAGACAC | SDM *k13* shield fwd | p7935 |
| **p6** | GTGTCTTTTTTAACCAgTTgATgAATGTAATTCTTAAATCACCTACCATATTTTC | SDM *k13* shield rev | p7936 |
| **p7** | GAGGTACCGAGCTCGAATTCGTAGCGAGAATGATTCTAATTC | *k13* CRISPR/Cas9 donor EcoRI In-Fusion fwd | p7942 |
| **p8** | CGAAAAGTGCCACCTGACGTCTGTTTATAACCATTAGATATATCAATATC | *k13* CRISPR/Cas9 donor AatII In-Fusion rev | p7943 |
| **p9** | AACATATGTTAAATATTTATTTCTC | CRISPR/Cas9 donor sequencing fwd | p282 |
| **p10** | AGGGTTATTGTCTCATGAGCGG | CRISPR/Cas9 donor sequencing rev | p283 |
| **p11** | AAGCACCGACTCGGTGCCAC | CRISPR/Cas9 gRNA sequencing rev | p35 |
| **p12** | GGGAATCTGGTGGTAACAGC | *k13* integration/sequencing primer fwd (5' end) to verify A212T | p6176 |
| **p13** | CGGAGTGACCAAATCTGGGA | *k13* integration/sequencing primer rev (3' end) to verify R539T/C580Y | p6175 |

*Shield refers to silent shield mutations introduced at the gRNA cut site to prevent cleavage of the recombinant locus following gene editing.

fwd, forward; rev, reverse; SDM, site-directed mutagenesis.

**Data S1. (separate excel file)**

Normalized metabolite peak intensities of DHA and OZ439 treated samples alongside untreated controls for Cam3.II isogenic lines harboring the following K13 genotypes; Wild-type, A212T, R539T and R539T+A212T.
